# EXO70D isoforms mediate selective autophagic degradation of Type-A ARR proteins to regulate cytokinin sensitivity

**DOI:** 10.1101/2020.02.07.938712

**Authors:** Atiako Kwame Acheampong, Carly Shanks, Chai-Yi Chang, G. Eric Schaller, Yasin Dagdas, Joseph J. Kieber

## Abstract

The phytohormone cytokinin influences many aspects of plant growth and development, several of which also involve the cellular process of autophagy, including leaf senescence, nutrient re-mobilization, and developmental transitions. The *Arabidopsis* type-A Response Regulators (type-A ARR) are negative regulators of cytokinin signaling that are transcriptionally induced in response to cytokinin. Here, we describe a mechanistic link between cytokinin signaling and autophagy, demonstrating that plants modulate cytokinin sensitivity through autophagic regulation of type-A ARR proteins. Type-A ARR proteins were degraded by autophagy in an AUTOPHAGY-RELATED (ATG)5-dependent manner. EXO70D family members interacted with Type-A ARR proteins, likely in a phosphorylation-dependent manner, and recruited them to autophagosomes via interaction with the core autophagy protein, ATG8. Consistently, loss-of-function *exo70D1,2,3* mutants compromised targeting of type-A ARRs to autophagic vesicles, have elevated levels of type-A ARR proteins, and are hyposensitive to cytokinin. Disruption of both type-A *ARRs* and *EXO70D1,2,3* compromised survival in carbon-deficient conditions, suggesting interaction between autophagy and cytokinin responsiveness in response to stress. These results indicate that the EXO70D proteins act as selective autophagy receptors to target type-A ARR cargos for autophagic degradation, demonstrating that cytokinin signaling can be modulated by selective autophagy.

## Introduction

Autophagy is a major catabolic pathway that maintains cellular homeostasis in response to intrinsic developmental changes and environmental cues. It mediates the degradation of protein complexes, misfolded and aggregated proteins, and damaged organelles by targeting proteins to proteases localized in the vacuole or lysosome, with a subsequent retro-transport of cellular building blocks back into the cytosol^1–3^. There are three main types of autophagy: macro-, micro- and chaperon-mediated autophagy^1, 4, 5^. Macro-autophagy (hereafter referred to simply as autophagy) is mediated by conserved AUTOPHAGY-RELATED GENEs (ATGs) that coordinate the *de novo* biogenesis of the autophagy organelle, the autophagosome^6–8^. Autophagosomes are double membrane vesicles that sequester various cargoes and ultimately deliver them to the lytic vacuole, resulting in their degradation and subsequent recycling. Although basal autophagy occurs in cells under steady-state conditions^9, 10^, it is enhanced in response to biotic and abiotic stresses^11–13^ and can result in selective or bulk (non-selective) degradation of proteins^4^. Typically, bulk autophagy randomly degrades cytosolic content, while selective autophagy requires unique receptors to target specific cargos for degradation^14, 15^. These receptors contain the unique ATG8-INTERACTING MOTIFs (AIM) or LIR-INTERACTING MOTIFs that allow them to interact with the ATG proteins on the autophagosomal membrane^16^. They also contain cargo-binding domains that mediate selective recruitment of specific cargo to the growing autophagosome. The presence of AIMs in various proteins suggests a link to autophagy. For example, autophagy has been linked to EXO70B1, a paralog of the EXO70 gene component of the exocyst complex, as it contains an AIM domain^17^. Consistent with this, EXO70B1 colocalizes with ATG8 proteins in autophagosomes and disruption of *EXO70B1* resulted in fewer vacuolar autophagic vesicles^17^. Multiple other EXO70 isoforms also contain AIM domains^18, 19^, suggesting that they may also act as receptors for the autophagic regulation of various cellular components.

The identification of receptors with their corresponding cargos has improved our understanding of the role of selective autophagy in regulating plant growth and development^15^. For example, in response to sulfur stress, Joka2, a tobacco member of the family of selective autophagy cargo receptors, triggers the autophagic degradation of the sulfur responsive protein, UPC9, by interacting with both UPC9 and ATG8f^20^. Selective autophagy also regulates rubisco degradation during leaf senescence^21^, brassinosteroid responses via targeting of BRI1-EMS SUPPRESSOR 1 (BES) by DOMINANT SUPPRESSOR OF KAR2 (DSK2)^22^, and ATG8-INTERACTING PROTEIN 1 (ATI1)-mediated turnover of plastid proteins^23^.

Autophagy has been linked to various phytohormones^24^, including cytokinins, which are *N*^6^- substituted adenine derivatives. Cytokinin and autophagy affect an overlapping set of plant developmental and physiological processes, including leaf senescence, nutrient remobilization, root meristem function, lateral root development, vascular development, and the response to biotic and abiotic stresses^24^. For example, cytokinin negatively regulates leaf senescence and plays a positive role in nitrogen uptake^25^. Disruption of autophagy in rice (via the *osatg7* mutant) results in early leaf senescence and compromised nitrogen reutilization^26^. Moreover, nitrogen remobilization between organs is reduced in autophagy-deficient mutants of *Arabidopsis* and maize^27, 28^. *ATG* genes are generally upregulated during leaf senescence^29^ and in response to biotic and abiotic stresses^12, 30^. Consistent with a link between cytokinin and autophagy, overexpression of a GFP-ATG8f fusion protein in *Arabidopsis* results in altered sensitivity to exogenous cytokinin, and cytokinin reduces the incorporation of this fusion protein into vacuolar-structures that are likely autophagic vesicles^31^. Further, transcriptomic analyses identified overlapping genes differentially expressed in *atg5-1* and cytokinin signaling mutants^32^. Despite these links, it is unclear how autophagy is integrated into cytokinin signaling.

The cytokinin signaling cascade in *Arabidopsis* is similar to bacterial two-component signaling systems (TCS) and begins with the binding of cytokinin to histidine kinase receptors, proceeds through a series of phospho-transfers, and culminates in the phosphorylation of conserved Asp residues in two classes of response regulators (ARRs): type-B and type-A ARRs^25, 33^. Type-B ARRs are DNA-binding transcription factors that mediate the transcriptional response to cytokinin, including the induction of type-A ARRs^34–36^. The *Arabidopsis* genome encode ten type-A ARRs which, unlike type-B ARRs, lack a DNA binding domain and act as negative regulators of cytokinin signaling^37^. Type-A ARR proteins generally have a short half-life and their levels are tightly regulated. Phosphorylation at the conserved Asp in response to cytokinin stabilizes a subset of the type-A ARRs^38^. Some type-A ARRs are degraded by the ubiquitin proteasome system^39, 40^, though there may be alternative ubiquitin-independent pathways regulating type-A ARRs turnover^38, 41^. For example, the nuclear-localized Periplasmic Degradation proteins (DEGs) selectively bind to and target ARR4 (but not other type-A ARRs) for ubiquitin-independent protein degradation^42^.

Here, we show that type-A ARRs are targeted for degradation by autophagy, at least in part through interaction with members of the EXO70D subfamily. *exo70D1,2,3* triple mutants have increased type-A ARR protein levels and reduced sensitivity to cytokinin. Our results suggest that these EXO70Ds target type-A ARRs for autophagic degradation, and thus mediate the interplay between cytokinin signaling and autophagy.

## Results

### Type-A ARR4 protein levels are regulated by autophagy

The level of type-A ARRs proteins plays a critical role in modulating the responsiveness to cytokinin in multiple developmental and physiological processes. Type-A ARRs levels are controlled both by regulation of their transcription by multiple inputs^43–46^ and by regulation of their protein stability, at least in part by phosphorylation of the conserved Asp residue^38–41^. Given the potential links between autophagy and cytokinin function, we tested if autophagy plays a role in the turnover of type-A ARR signaling elements. Inhibiting autophagy in transgenic *Arabidopsis* seedlings with autophagy inhibitor, concanamycin A (ConcA), resulted in the rapid accumulation of various type-A ARR:CFP/eGFP fusion proteins (ARR3, ARR4, ARR5, ARR7 and ARR16) (Fig. 1a; Supplementary Fig. 1a), but did not affect type-A *ARR* transcript levels (Supplementary Fig. 1b). This effect of ConcA on type-A ARR proteins was independent of the epitope tag or location as N-terminally tagged ARR7 (myc:ARR7) also accumulated in response to ConcA (Supplementary Fig. 1c). Consistent with the type-A ARRs being degraded by autophagy, treatment with ConcA induced the accumulation of multiple type-A ARR:CFP fusion proteins into vesicles, which likely correspond to autophagic vesicles (Supplementary Fig. 1d).

**Figure 1:**
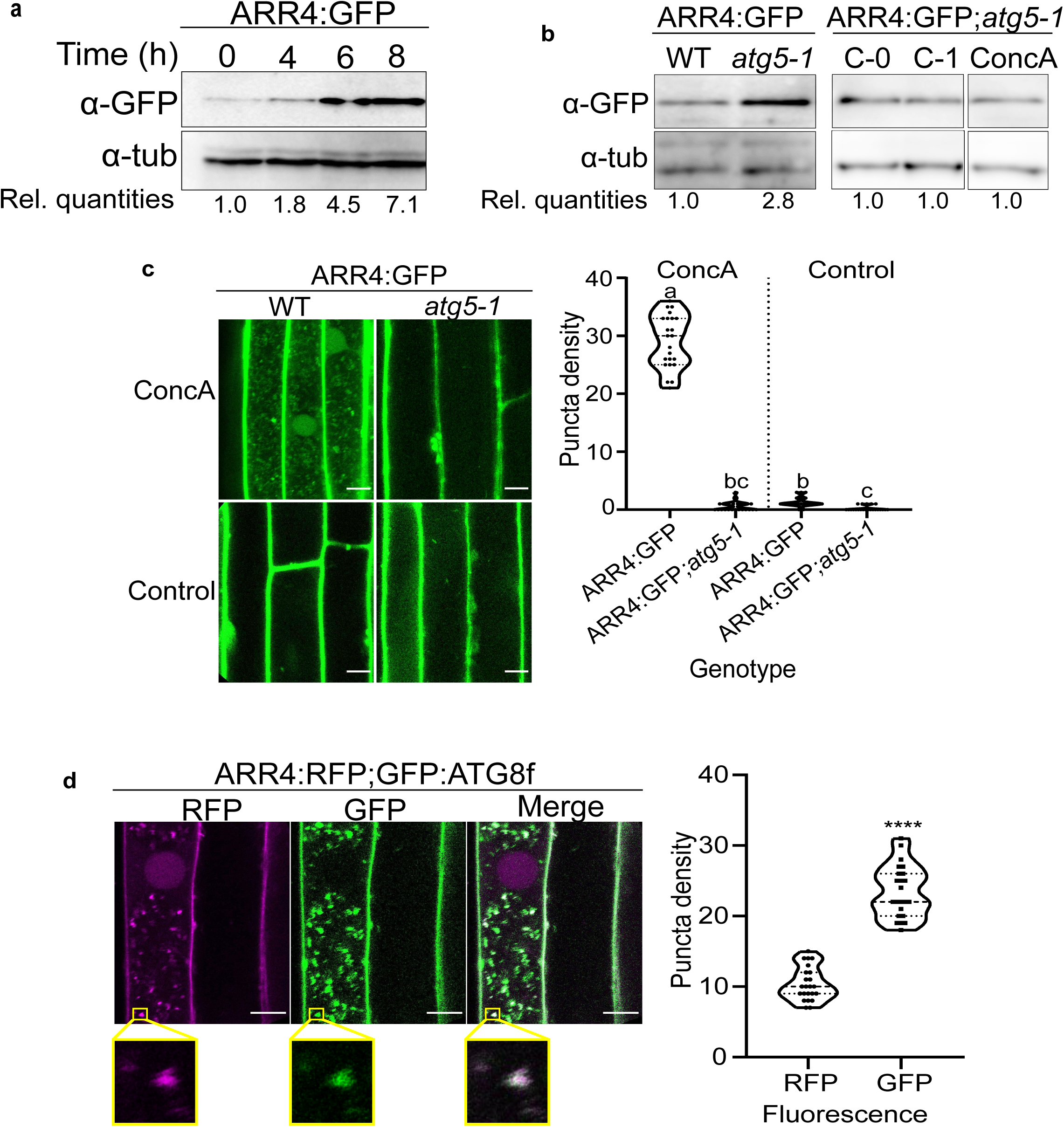
ARR4 is degraded by the autophagy pathway in *Arabidopsis* roots. **(a)** Response of root-expressed ARR4 to concanamycin A (ConcA), the inhibitor of autophagy. Seedlings carrying *pUBQ10::ARR4:GFP* constructs were treated for 8 h in sucrose-deficient liquid MS media supplemented with 1 µM ConcA and transferred to dark to induce carbon starvation. ARR4 protein quantities in the roots were analyzed by immunoblot assays using anti-GFP (α-GFP) antibodies. Anti-tubulin (α-tub) served as loading control. Rel. quantities represent ratio of intensity of α-GFP to α-tub band relative to ratio at 0 h. **(b)** Effect of disruption of autophagosome-forming *ATG5* gene on the levels of ectopically-expressed ARR4 proteins. *pUBQ10::ARR4:GFP-*expressing plants described in **(a)** was crossed to the *atg5-1*^1,2^. The progeny of the cross and the wild-type plants were grown on normal MS media and in light for nine days, and for additional 1 day under carbon starvation conditions. Seedlings were either untreated (C-0) or treated with mock control (C-1) and ConcA (ConcA) as described above. ARR4:GFP protein in the roots was analyzed by immunoblot assay. Rel. quantities represent ratio of intensity of α-GFP to α-tub band relative to ratio of WT band. **(c)** Representative confocal micrograph of the root elongation zone of five-day old *Arabidopsis* seedlings described in **(1b)**. The light grown seedlings were treated with ConcA or DMSO as control and exposed to carbon starvation for 18 h prior to imaging. Scale bar =10 µm. **(Right)** Quantification of the number of GFP-containing vacuolar puncta. Data represents the average number of puncta per 100 µm^2^ of vacuole area, n=23. Data was analyzed with one-way ANOVA followed by Tukey-Kramer multiple comparison analyses; *p*<0.05. Different letters represent statistically different means. **(d)** Intracellular colocalization of RFP signal resulting from ARR4:RFP with the GFP signal from GFP:ATG8f. Seedling co-expressing both constructs were grown for 5 days in MS medium followed by 18 h treatment with ConcA under conditions of carbon starvation. Scale bar=10 µm. The magnified area enclosed by the yellow box indicates section of vacuole showing colocalization of ARR4:RFP and GFP:ATG8f-containing vesicles. Graph represents mean (n=23) number of RFP- and GFP-containing puncta per 50 µm^2^ of vacuole area. **** indicate statistical differences at *p*<0.0001 using the Student *t*-test **(Right)**.

To further explore the role of autophagy in type-A ARR turnover, we focused on a transgenic line expressing an ARR4:eGFP transgenic protein. Similar to the effect of ConcA, genetic disruption of autophagy (via the *atg5* mutation) resulted in elevated ARR4:GFP protein levels, without affecting ARR4 transcript levels (Fig. 1b and Supplementary Fig. 1e). Further, the accumulation of ARR4:GFP into vesicles was nearly eliminated in *atg5-1* mutants (Fig. 1c), indicating that these vesicles are indeed autophagic vesicles. The level of ARR4:GFP protein in the *atg5-1* background was unaffected by ConcA treatment, consistent with the increased level of ARR4 in response to ConcA being the result of disrupted autophagy (Fig. 1b).

To confirm that the ConcA-dependent ARR4-containing vesicles are autophagic vesicles, we analyzed the intracellular localization of ARR4:RFP- and GFP:ATG8f-containing vesicles in roots of stable transgenic *Arabidopsis* plants. Overall, there were more than twice as many GFP:ATG8f vesicles as ARR4:RFP vesicles (Fig. 1d). Post-thresholded Manders’ Colocalization Coefficients^47^ values indicate that 81% of the ARR4:RFP vesicles overlapped with GFP:ATG8f vesicles, while 52% of GFP:ATG8f associate with compartments containing ARR4:RFP. This analysis indicates that the ARR4:RFP signal significantly colocalized with GFP:ATG8f, consistent with ARR4 being targeted to a subset of autophagic vesicles.

### Type-A ARRs interact with the EXO70D proteins

To explore the mechanism of autophagic regulation of type-A ARRs and to identify potential receptors involved in targeting type-A ARR proteins to autophagic vesicles, we screened for proteins that interact with an activated form of type-A ARR protein. Previous studies indicate that mutating the aspartic acid residue (D) that is the target of phosphorylation in the receiver domain of ARRs to a glutamic acid (E) partially mimics the activated, phosphorylated form of the protein. Using an ARR5^D87E^ bait in a yeast two-hybrid screen, we identified three independent preys that corresponded to the *EXO70D3* (AT3G14090) gene. EXO70D3 belongs to the 23-member QR-motif-containing *Arabidopsis* EXO70 gene family^48, 49^ and is most closely related to two other paralogs, EXO70D1 (AT1G72470) and EXO70D2 (AT1G54090) (Supplementary Fig. 2a). EXO70s are subunits of the octomeric exocyst complex, which is involved in exocytosis and other cellular trafficking processes. Recent studies have shown that EXO70 paralogs could also play roles in autophagy^17, 50, 51^. EXO70D3 interacted with ARR5^D87E^, but not with a wild-type ARR5 or ARR5^D87A^ bait in a yeast two-hybrid assay (Fig. 2a), suggesting it may interact with ARR5 in a phospho-Asp-dependent manner. We determined the interactions among the EXO70D isoforms and multiple type-A ARRs using a yeast two-hybrid assay (Fig. 2a). The three EXO70D paralogs interacted differentially with a subset of type-A ARRs, generally preferentially with their phosphomimic forms. The D→E version of ARR4 (ARR4^D94E^) interacted with all three EXO70D paralogs, but neither the wild type nor the D→A version of ARR4 interacted appreciably with any EXO70D isoform in this assay. All three versions of ARR5 (ARR5, ARR5^D87A^ and ARR5^D87E^) interacted strongly with EXO70D1, but only ARR5^D87E^ interacted with EXO70D2 and EXO70D3. The D→E version of ARR7 (ARR7^D85E^) interacted with EXO70D1, but EXO70D2 and EXO70D3 did not appreciably interact with any version of ARR7. The D→E version of ARR16 (ARR16^D93E^) interacted very weakly with EXO70D1, but not with EXO70D2 nor EXO70D3. These results suggest that all three paralogs of EXO70D may play a role in regulating type-A ARR function, with some degree of specificity.

**Figure 2:**
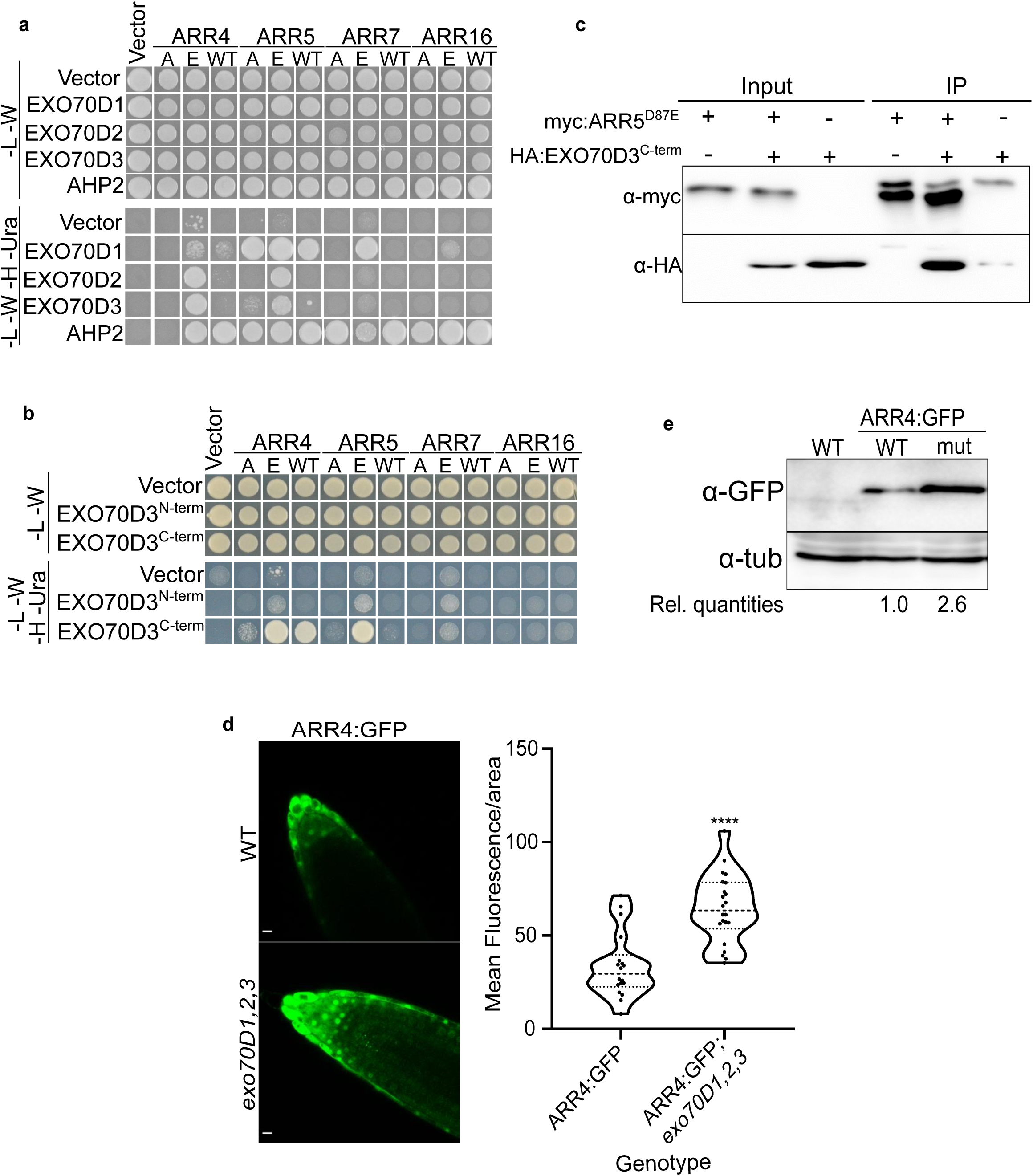
Interaction between EXO70Ds and type-A ARRs results in the destabilization of the type-A ARRs. **(a)** Yeast two-hybrid (Y2H) assay showing pairwise interactions between members of the EXO70D sub-clade and representative type-A ARRs. The EXO70D paralogs were cloned into the Gal4 DNA Activation domain prey vector while the type-A ARRs (WT) with their respective phosphor-dead (A) and phosphor-mimic (E) mutations were cloned as baits in Gal4 DNA binding domain vectors. Interactions were accessed on -Leu-Trp-His-Ura (-L -W -H -Ura) quadruple dropout media. The -Leu-Trp (-L -W) served as control media. The empty bait and prey vectors were used as negative controls, whereas the interactions between the type-A ARRs and *Arabidopsis* Histidine Phosphotransfer Protein 2 (AHP2) served as positive control. **(b)** Y2H assay of interactions of N-terminal (EXO70D3^N-term^) or C-terminal (EXO70D3^C-term^) domains of EXO70D3 with the representative type-A ARRs described in **(2a)**. See Supplementary Fig. 3a for details on N-term and C-term. Interaction assays is similar to **(2a)**. **(c)** EXO70D3^C-term^ interacts with phosphor-mimic mutants of ARR5 (ARR5^D87E^) in co-immunoprecipitation assay. Leaves of *Nicotiana benthamiana* were infiltrated with plasmids expressing myc:EXO70D3^C-term^, or HA:ARR5^D87E^. The input extracts and the myc-immunoprecipitated proteins (IP) were analyzed by immunoblotting with anti-HA and anti-myc. **(d)** Ectopically-expressed ARR4:GFP accumulates in roots tips of *exo70D1,2,3* triple loss-of-function mutant. Confocal microscopy images indicating the expression of pUBQ10::ARR4:GFP in wild type (top panel) and *exo70D1,2,3* mutant (bottom panel) plants. Scale bars = 10 µm. **(Right)** Images are quantified by measuring signal intensity of individual nuclei using FIJI software, after background normalization. The graph represents the average signal measurement from 22 images of each genotype. Data was analyzed by unpaired Student’s t-test. **** indicate statistical difference at *p*<0.001 (4.025e-006). **(e)** Anti-GFP immunoblot analyses of roots of 10-day old *Arabidopsis* seedlings, transgenic seedlings expressing *pUBQ10::ARR4:GFP* in wild type (WT) or *exo70D1,2,3* (mut) described in **(2d)**. Rel. quantities represent ratio of intensity of α-GFP to α-tub band relative to ratio of bands from WT (ARR4:GFP).

We further confirmed the phospho-Asp-dependent interaction of ARR5 and EXO70D3 *in planta* using bimolecular fluorescence complementation (BiFC) (Supplementary Fig. 2b). Transient co-expression of cYFP:ARR5^D87E^ and nYFP:EXO70D3 in leaves of *Nicotiana benthamiana* resulted in stronger fluorescence from the reconstituted YFP as compared to co-expressed cYFP:ARR5^D87A^ and nYFP:EXO70D3. These BiFC interactions occurred in the cytosol and are consistent with the localization of a subset of type-A ARRs partially in the cytosol of onion skin epidermal cells^52–54^ and the exclusive localization of UBQ10::EXO70D3:GFP in the cytosol of *Arabidopsis* roots (Supplementary Fig. 2c) and in transfected *Arabidopsis* protoplasts^55^.

Analysis of EXO70D3 using the iLIR database^56^ revealed the presence of an AIM ^56^ (xLIR: L^232^-V^237^) in the N-terminal domain (Supplementary Fig. 3a), suggesting it may interact with ATG8 proteins and act as a receptor to mediate the targeting of type-A ARRs to autophagosomes. Both EXO70D1 and EXO70D2 also contain EXO70 domains (EXO70D1: R^247^-D^617^ and EXO70D2: R^239^-D^607^) and putative AIMs (EXO70D1: WxxL motif: L^237^-L^242^, and “anchor”: D^90^-I^95^; EXO70D2: xLIR: L^229^-V^234^) (Supplementary Fig. 3b), which are suggestive of roles as autophagy receptors.

To identify the domain of EXO70D3 that interacts with type-A ARRs, we performed pairwise yeast-2 hybrid assays using the N-terminus (EXO70D3^N-term^; amino acid: 1-298; includes the AIM domain) or C-terminus (EXO70D3^C-term^; amino acid: 299-623) (Supplementary Fig. 3a) of EXO70D3 with wild-type or phospho-mutant forms of type-A ARR paralogs (Fig. 2b). While an EXO70D3^N-term^ bait did not interact with type-A ARRs, an EXO70D3^C-term^ bait interacted most strongly with the phosphomimic versions of ARR4 and ARR5, but weakly or not at all with ARR7 and ARR16, consistent with the results observed using the full-length protein. We confirmed the interaction of ARR5 and EXO70D3 *in vivo* using co-immunoprecipitation (co-IP) with ARR5^D87E^ and EXO70D3^C-term^ (Fig. 2c). We conclude that the C-terminal domain of EXO70Ds interacts preferentially with the phosphorylated form of multiple type-A ARRs.

### EXO70D3 regulates Type-A ARR protein levels

We examined the consequence of the interaction of EXO70Ds and type-A ARRs by analyzing the effect of modulating EXO70D function on type-A ARR protein levels. When transiently co-expressed in *N. benthamiana* leaves, EXO70D3 reduced ARR5^D87E^ protein levels (Supplementary Fig. 3c). Co-expression with a negative control plasmid had no effect on ARR levels. Although they did not interact in our yeast-two hybrid assay, EXO70D3 reduced ARR5^WT^ proteins in a concentration-dependent manner when co-infiltrated in *N. benthamiana* leaves (Supplementary Fig. 3d), suggesting that these proteins may interact *in planta*, possibly reflecting a weak interaction insufficient for the yeast two-hybrid assay or *in planta* phosphorylation of ARR5^WT^ in this system. We examined the effect of separating the N- and C-terminal domains of EXO70D3 on degradation of ARR5 *in planta*. Co-expressing either the EXO70D3^N-term^ or EXO70D3^C-term^ alone with ARR5^D87E^ did not significantly affect its stability (Supplementary Fig. 3e), indicating that both the type-A ARR-interacting (EXO70D3^C-term^) and AIM-containing domains (EXO70D3^N-term^) are required for effective destabilization of ARR5. This is consistent with EXO70D3 acting as a receptor that recruits type-A ARR cargos for autophagic degradation, requiring both type-A ARR and ATG8 (i.e. AIM motif) binding domains.

All three paralogs EXO70Ds are expressed in *Arabidopsis* roots (Supplementary Fig. 3f), and thus we examined the effect of disruption of the EXO70Ds on type-A ARR stability in this tissue. Disruption of *EXO70D1*, *EXO70D2* and *EXO70D3* resulted in a significant increase in fluorescence in roots of an *Arabidopsis* line expressing GFP-tagged ARR4 from a *UBQ10* promoter (Fig. 2d and Supplementary Fig. 3g). We confirmed this result using an independent line carrying a *UBQ10::ARR4:RFP* transgene in a wild type and *exo70D1,2,3* triple mutant background (Supplementary Fig. 3h). Further, immunoblotting revealed a ∼2.5 increase in ARR4:GFP levels in *exo70D1,2,3* as compared to a wild-type background (Fig. 2e).

### EXO70D3 promotes targeting of ARR4 to autophagic vesicles

The results described above suggest that the EXO70D paralogs play a role in the autophagic turnover of at least a subset of type-A ARRs^55,57–59^. Other members of EXO70 gene family have been linked to autophagy and the formation of autophagosomes^17, 19, 60^. Thus, we tested the hypothesis that the EXO70Ds target type-A ARRs to autophagic vesicles. In the roots of stable transgenic *Arabidopsis* plants, disrupting all three members of the EXO70D gene family resulted in a 50% reduction in the number of vacuolar, ConcA-dependent ARR4:GFP-containing vesicles (Fig. 3a). Consistent with this, the increase in ARR4:GFP protein levels in response to ConcA was largely dependent on functional *EXO70D* genes (Fig. 3b). These results suggest that EXO70Ds regulate ARR4 protein levels via autophagy.

**Figure 3:**
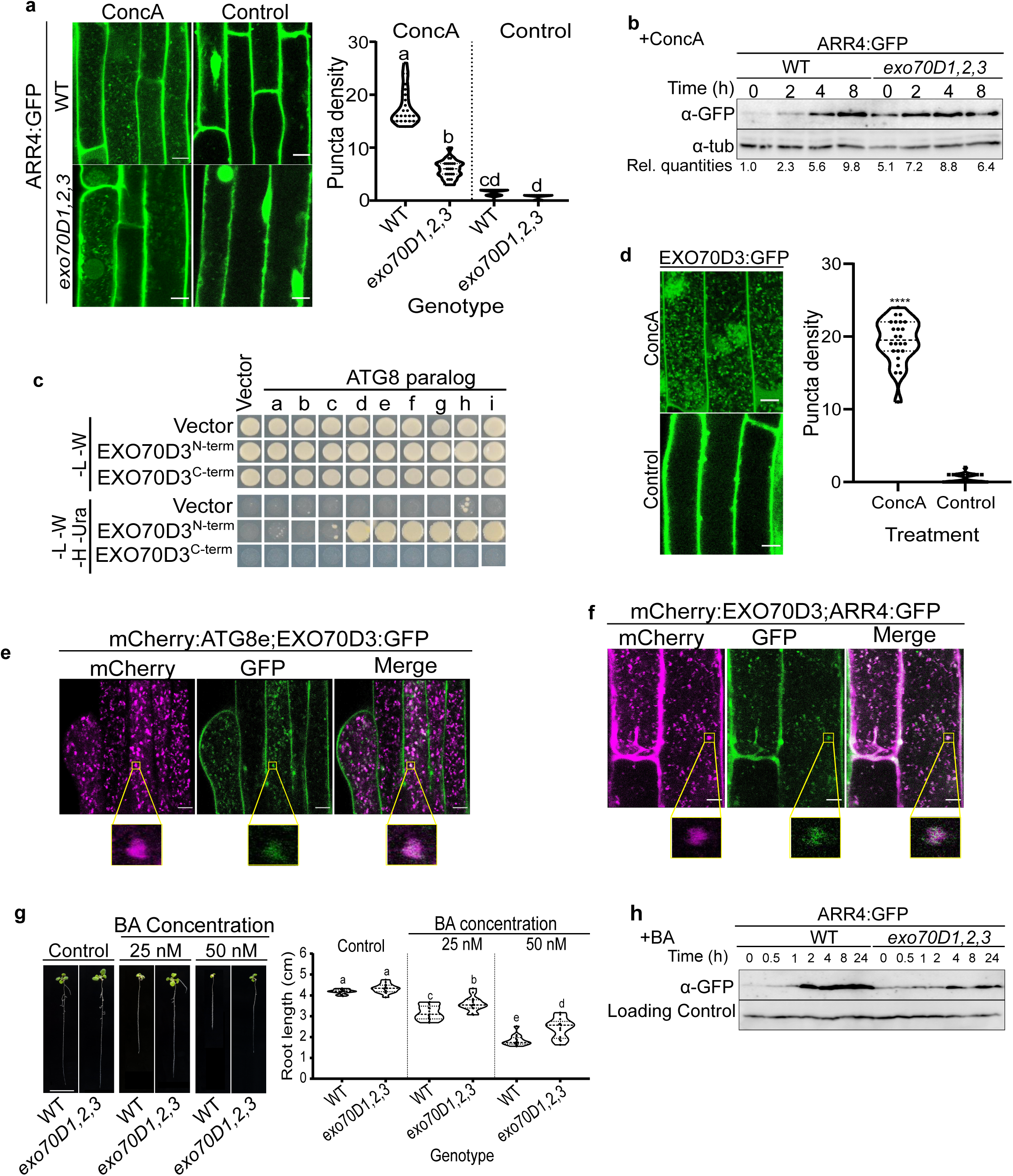
EXO70Ds mediate cytokinin responses by recruiting type-A ARRs to the ATG8-tagged autophagy machinery for degradation. **(a)** Representative confocal micrographs of the root elongation zones of ectopically-expressed ARR4:GFP in wild type and *exo70D1,2,3* mutants. Seedlings were grown for 5 days on MS plates, followed by incubation with 1 µM ConcA or DMSO (as control) under carbon starvation conditions for 18 h. Scale bar =10 µm. Quantification of the number of GFP-containing puncta is as described in Figure 1c. Data (n=23) was analyzed with one-way ANOVA followed by Tukey-Kramer multiple comparison analyses; *p*<0.05. Different letters represent values that are statistically different at the indicated *p*-value. **(b)** EXO70D-dependent regulation of ARR4 proteins in response to the autophagy inhibitor, ConcA. Ten-day old seedlings were treated with 1 µM ConcA followed by immunoblot assays using anti-GFP. α-tub was loading control. Rel. quantities represent ratio of intensity of α-GFP to α-tub band relative to ratio of bands from untreated WT (ARR4:GFP; 0h). **(c)** Yeast two-hybrid interaction of ATG8 paralogs with N-terminal (EXO70D3^N-term^) or C-terminal (EXO70D3^C-term^) domains of EXO70D3. Interactions were analyzed on Leu-Trp-His-Ura (-L -W -H -Ura) quadruple dropout media, with -Leu-Trp (-L -W) double dropout serving as control media. **(d)** Representative confocal microscopy image of the root elongation zones of 5-day old *Arabidopsis* seedlings expressing EXO70D3:GFP. ConcA or DMSO (as control) treatment is same as described in **(3a)**. Scale bar =10 µm. Quantification of the number of EXO70D3:GFP-containing puncta. Data, representing 24 independent images, was analyzed with Students’ *t*-test; *p*<0.05. Asteriks (****) represent values that are statistically different at *p*<5.066×10^-0321^. **(e)** Colocalization of EXO70D3:GFP and mCherry:ATG8e in roots of *Arabidopsis* seedlings carrying expression plasmids for both genes. Prior to imaging, seedlings were incubated for 18 h in the dark in sucrose-deficient media supplemented with 1 µM ConcA. **(f)** Colocalization of mCherry:EXO70D3 and ARR4:GFP in elongation zone of 5-day old *Arabidopsis* seedlings ectopically-expressing both genes. Seedling treatment and imaging is as described for **(3e)**. The magnified areas enclosed by the yellow boxes in (e) and (f) show colocalization of vesicles containing mCherry:ATG8e and EXO70D3:GFP, and Cherry:EXO70D3 and ARR4:GFP, respectively. **(g)** Root elongation assay showing response of roots of *exo70D1,2,3* and wild-type seedlings to exogenous cytokinin. Seedlings were grown on MS media for 4 days, and transferred to MS media supplemented with BA (cytokinin) or NaOH (control) for 5 days. Scale bar = 1 cm. Plants were imaged and root length measured using FIJI software. Average length (n≥13) was analyzed using one-way ANOVA followed by Tukey-Kramer multiple comparison analyses at *p*<0.05. Different letters represent statistically different means. The assay was conducted three times with similar results. **(h)** Effect of BA treatment on ARR4:GFP protein levels in wild-type and *exo70D1,2,3* mutant seedlings. Proteins extracted from 10-day old seedlings following treatment with cytokinin (BA) for up to 24 h, were analyzed by immunoblot assay using anti-GFP. Anti-tubulin (α-tub) served as loading control.

If the EXO70D proteins act as autophagy receptors, they should interact with various ATG8 isoforms and should themselves be targeted to autophagic vesicles. Indeed, EXO70D3^N-term^ interacted with multiple members of the *ATG8* gene family in a yeast-two hybrid assay (Fig. 3c), consistent with the presence of AIMs in this domain. Furthermore, we observed ConcA-dependent EXO70D3:GFP-containing vesicles in *Arabidopsis* roots (Fig. 3d). In *Arabidopsis* seedlings co-expressing EXO70D3:GFP and mCherry:ATG8e, these ConcA-dependent EXO70D3-containing puncta colocalized with mCherry:ATG8e-tagged puncta (Fig. 3e). To test whether these EXO70D3-containing puncta are related to the ARR4-containing vesicles, we analyzed the localization of these proteins in stable transgenic *Arabidopsis* plants. The overlap between the mCherry and GFP-containing puncta indicates that both EXO70D3 and ARR4 are recruited into the autophagic vesicles (Fig. 3f). These observations are consistent with the EXO70Ds acting as receptors for the selective autophagy of ARR4 by recruiting the ARR4 to the autophagosome via the interaction between EXO70D3 and ATG8.

### Perturbation of EXO70Ds alters cytokinin response

In agreement with previous reports^48^, we did not observe any substantial morphological changes in ten-day-old single (*exo70D1*, *exo70D2* or *exo70D3),* double (*exo70D1,2, exo70D1,3, exo70D2,3*) or triple (*exo70D1,2,3*) mutants as compared to wild type (Fig. 3g & Supplementary Fig. 4). Six-week old single, double and triple mutant adult plants were also comparable to the wild type (Supplementary Fig. 4).

We examined the link between EXO70D3 function and cytokinin. The *EXO70D* genes are not transcriptionally regulated by cytokinin (Supplementary Fig. 5a). Prior studies revealed that overexpression of type-A ARRs results in hyposensitivity of primary roots to cytokinin^38, 41^, and as type-A ARR protein accumulates in the *exo70D1,2,3* triple mutant (Fig. 2d & Fig. 2e), we examined if the *exo70D1,2,3* mutant is altered in the response to exogenous cytokinin. The *exo70D1,2,3* mutants showed reduced sensitivity to low doses of exogenous cytokinin in a root elongation assay (Fig. 3g). The single (*exo70D1*, *exo70D2*, *exo70D3*) and double (*exo70D1,2*, *exo70D1,3*, *exo70D2,3*) mutants did not exhibit altered cytokinin responsiveness (Supplementary Fig. 5b & 5c). These results demonstrate that members of the EXO70D gene family act redundantly to positively regulate cytokinin responsiveness.

To investigate the effect of disruption of EXO70Ds on the spatial pattern of the cytokinin response, we examined the expression of a TCSn::GFP cytokinin reporter in roots of wild type and *exo70D1,2,3* mutants grown in the presence of cytokinin. In the absence of exogenous cytokinin, TCSn::GFP expression was reduced in the *exo70D1,2,3* mutant as compared to wild-type roots (Supplementary Fig. 5d). When grown in the presence of exogenous cytokinin, TCSn::GFP expression in the *exo70D1,2,3* mutant was significantly lower compared to wild type. Taken together, these results suggest that disruption of the *EXO70D*s results in reduced responsiveness to endogenous and exogenous cytokinin, likely as a consequence of elevated type-A ARR protein levels.

Cytokinin was previously shown to stabilize a subset of type-A ARR proteins, likely at least in part by blocking targeting to the 26S proteasome^38^. To explore the interaction of autophagy with this process, we analyzed the effect of cytokinin on type-A ARR protein levels in an *exo70D1,2,3* triple mutant. Interestingly, though exogenous cytokinin resulted in accumulation of ARR4:GFP protein in the roots of both wild-type and *exo70D1,2,3* seedlings, the rate of accumulation in the triple mutant was slower than in the wild type (Fig. 3h). This suggests that disrupting EXO70D-mediated autophagy may up-regulate other cellular mechanisms modulating type-A ARR stability, such as ubiquitin-mediated degradation via the 26S proteasome.

### EXO70Ds and type-A ARRs enhance response to fixed-carbon starvation

Mutants that disrupt some *ATG* genes or receptors of autophagy are hypersensitive to or have reduced survival in nitrogen or carbon starvation assays^17, 22, 61^. Thus, we assayed the response of the single, double and triple mutants of *EXO70D*s to carbon starvation. Consistent with a role in autophagy, *exo70D1,2,3* is hypersensitive to carbon starvation; After seven days of dark treatment, the survival rate of *exo70D1,2,3* was 75% compared to 100% for wild-type plants. After eleven days of dark treatment, the survival decreased to 50% for *exo70D1,2,3,* while it was greater than 80% for wild-type plants (Fig. 4a). Responses of single mutants were generally comparable to that of wild-type plants after seven and eleven days of dark treatment (Supplementary Fig. 6a). Among the double mutants, eleven days of dark treatment resulted in 50% and 80% survival for *exo70D1,3* and *exo70D2,3* respectively, compared to about 90% for *exo70D1,2* and wild-type plants (Supplementary Fig. 6b). Overall, these results suggest that members of the *EXO70D* gene family redundantly function in regulating autophagic responses during carbon starvation.

**Figure 4:**
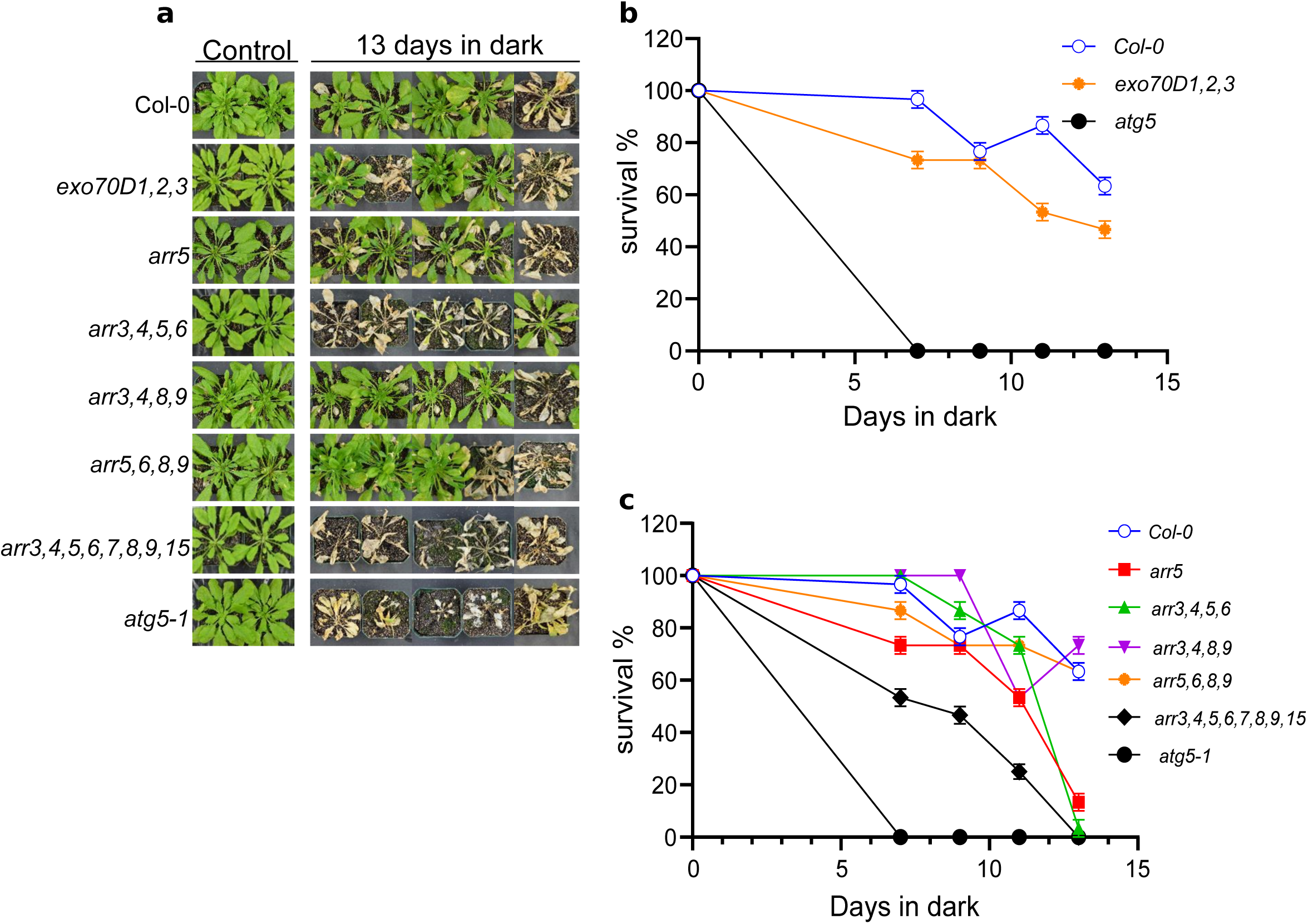
EXO70Ds and type-A ARRs modulate plant response to fixed-carbon starvation. Seedlings were grown for six weeks on potted soil under short day conditions of 16/8 h day/night at 22 °C. Pots were transferred to dark for 7, 9, 11, 13 days, and recovered in the light for 7 days. **(a)** Representative images of *exo70D1,2,3* triple loss-of-function mutants, *arr5-1* and some high-order type-A ARR loss-of-function mutants (*arr3,4,5,6*; *arr3,4,8,9*; *arr5,6,8,9*; *arr3,4,5,6,7,8,9,15*) following 13-day-dark treatment. Col-0 and *atg5-1* served as wild-type and autophagy deficient controls, respectively. Graphs represent quantification of survival of *exo70D1,2,3* **(b)**, and various mutants of type-A ARR **(c)** in response to carbon starvation. Survival was estimated as the percentage of plants with new leaves after the dark treatment. Values represent mean ± SEM percentage survival of 3 biological replicates. Each biological replicate consisted of 8 plants per genotype per treatment.

We tested if the type-A ARRs also play a role in the response to carbon starvation by examining the phenotypes of high-order type-A ARR loss-of-function mutants in fixed-carbon starvation survival assays. Compared to wild-type plants, the type-A ARR higher order mutants were significantly more sensitive to carbon starvation (Fig. 4b), with the *arr3,4,5,6,7,8,9,15* octuple mutant displaying the strongest phenotype, suggesting at least partial functional redundancy among the type-A ARRs in this response.

## Discussion

Autophagy and cytokinin regulate overlapping plant growth and developmental processes, including senescence, root and vascular development, and nutrient remobilization^4, 5, 13, 25, 33^. Further links between autophagy and cytokinin include the observations that *atg7* mutants in rice have decreased levels of cytokinins^62^ and over-expression of *ATG8* enhances cytokinin perception in *Arabidopsis*^31^. Here, we demonstrate that plants modulate responses to cytokinin, at least in part, via the autophagic regulation of type-A ARRs. This novel input into cytokinin function is mediated by members of the EXO70D subclade of the EXO70 gene family, which act as receptors to recruit type-A ARRs to the autophagosome for subsequent degradation. This conclusion is based on: 1) The interaction of EXO70Ds with a subset of type-A ARRs and with ATG8 paralogs; 2) The colocalization of EXO70D3, type-A ARRs, and ATG8 in vacuolar vesicles; 3) The elevated level of type-A ARR proteins in *atg5* and *exo70D3* mutants; and 4) The reduction in the number of ConcA-dependent ARR containing vesicles in *exo70D1,2,3* mutants. The elevated levels of type-A ARR proteins in *atg5* and *exo70D1,2,3* mutants result in hyposensitivity to cytokinin, similar to prior studies on transgenic lines overexpressing type-A ARRs^38, 41^, which, like the *exo70D1,2,3* triple mutant, are largely aphenotypic when grown under normal lab conditions. Thus, autophagy regulates the sensitivity to cytokinin, at least in part via EXO70Ds. The residual trafficking of ARR4 into autophagic vesicles in *exo70D1,2,3* mutants may reflect additional receptors involved in the selective autophagy of type-A ARRs. The ARR-EXO70D-ATG8 interactions and the responses to cytokinin and carbon deficient conditions demonstrated in this study provide a molecular mechanism linking autophagy and cytokinin responses.

EXO70 genes are highly expanded in *Arabidopsis* and other land plants^48, 49, 63^, are expressed in specific tissue types^64^, and regulate diverse physiological processes^65^. For example, Exo70A1 is implicated in the recycling of PINs at the plasma membrane and polar growth^48, 66^, development of tracheary elements^67^, cell cycling^68^, Casparian strip development^69^, and cell plate formation^70^. EXO70C2 regulates pollen tube growth^71^ and Exo70B1 and Exo70H1 regulate pathogen defense responses^72, 73^. Here we demonstrate a novel role for the EXO70D clade of this expanded gene family. While the results presented here suggest that the phosphorylation of a subset of type-A ARRs, which occurs in response to cytokinin, increases their targeting to autophagic degradation, prior studies indicated that cytokinin stabilizes a subset of type-A ARRs proteins, also likely through phosphorylation of the conserved Asp in their receiver domain^38^. Further, while our findings show that type-A ARR proteins are regulated by autophagy, other reports indicate that the 26S proteasome also plays a role in the turnover of type-A ARRs^40, 41, 74^. Similar proteasome-dependent and -independent regulator pathways have been reported for BRI1-EMS SUPPRESSOR 1^22^ and for a number of ABA signaling components^75^.

We propose a model (Fig. 5) in which the unphosphorylated type-A ARRs are targeted for degradation by the 26S proteasome. In response to cytokinin, type-A transcript levels rise and the type-A ARR proteins become phosphorylated, reducing their targeting to the 26S proteasome, but ultimately increasing their degradation by autophagy. The relative contributions of these two mechanisms to type-A ARR protein turnover is likely influenced by tissue/cell type, the particular type-A ARR isoform involved, and other regulatory inputs. Together, these two mechanisms likely complement each other to optimally tune cytokinin responsiveness in response to various developmental and environmental cues.

**Figure 5:**
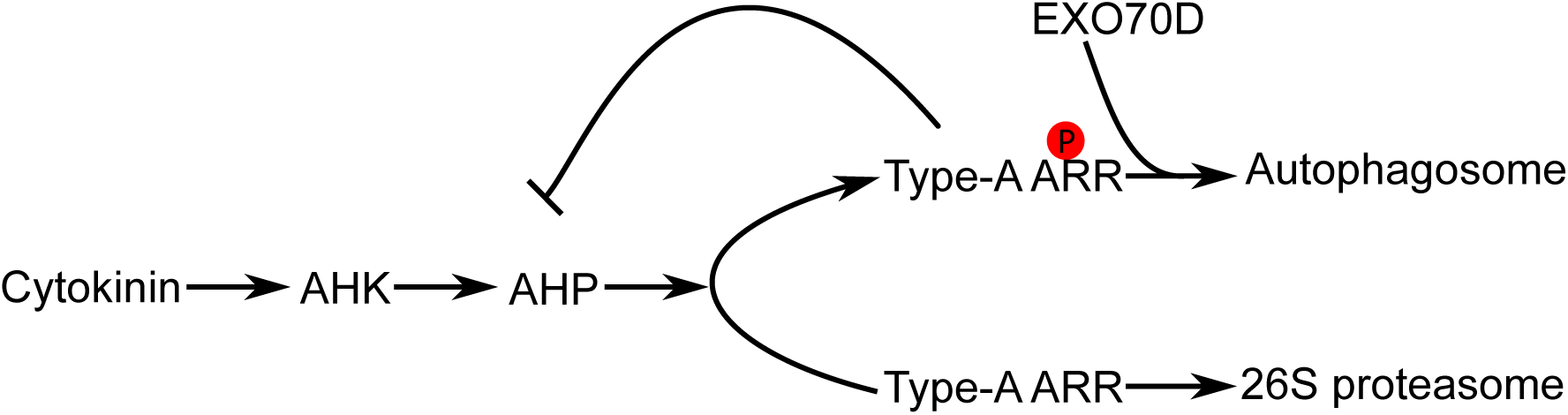
Model of autophagic and proteasome regulation of type-A ARRs. In the presence of cytokinin, type-A ARRs are phosphorylated on conserved Aspartate in the receiver domain. Phosphorylated type-A ARRs are more stable and negatively regulate cytokinin responses. To regulate this constitutive type-A ARR action, EXO70Ds recruit the phosphorylated type-A ARR to the autophagosome by interacting with ATG8 isoforms. This EXO70D-mediated autophagic mechanism presents a more rapid degradation pathway for type-A ARRs. In the absence of cytokinin, unphosphorylated type-A ARRs are ubiquitinated and shuttled to the 26S proteasome for degradation. This 26S proteasome pathway is probably activated for a prolong degradation of type-A ARRs. Black arrow: phosphate transfer; Green line: protein-protein interaction; Purple Arrow: protein degradation.

Although *exo70D1,2,3* is morphologically indistinguishable from wild-type plants under normal growing conditions, it is significantly compromised in survival under carbon deficient conditions. Consistent with this, mutations in autophagy receptors, such as *exo70B1* and *dsk2,* result in reduced survival in nutrient deficient conditions^22, 76^. The observation that disruption of all three *EXO70D*s resulted in a less severe effect as compared to mutations disrupting bulk autophagy regulators (i.e. *ATG5*) is consistent with a role for EXO70Ds acting as receptors for selective autophagy.

The observation that disruption of type-A ARRs significantly compromises plant survival under carbon-limiting conditions suggests there is feedback between autophagy and cytokinin signaling. As cytokinin regulation has pleiotropic effect on plant development, including senescence and nutrient partitioning^25, 33^, it is possible that disrupting a central cytokinin signaling component such as type-A ARRs may significantly affect responses to carbon limitation indirectly, rather than by directly modulating autophagy. For example, cytokinin promotes localized sink activity^77^, and thus disruption of type-A ARRs may affect nutrient partitioning, which in turn would likely impact survival in response to carbon starvation. Cytokinin also plays a key role in regulating leaf senescence^78^, which may impact the response to carbon starvation. Alternatively, it is possible that cytokinin signaling has a direct effect on autophagy via an as yet unidentified mechanism.

In conclusion, our findings have uncovered a novel regulatory pathway by which type-A ARR are recruited by EXO70Ds to the autophagosome for degradation. Previous reports indicate that subset of type-A ARRs are regulated by the 26S proteasome^39, 40^ and by a ubiquitin-independent protein pathway involving the DEG9 protein^42^. As type-A ARRs are primary cytokinin response genes and negatively regulate the cytokinin signaling pathway, it is perhaps not surprising that they are regulated through multiple pathways. The relative contributions of these regulatory mechanisms, the cellular and environmental conditions under which they are triggered, the cell types in which they predominantly occur, and the specific type-A ARR isoforms that are regulated by each mechanism remain to be determined. As type-A and type-B ARRs both possess receiver domains that are phosphorylated by AHPs^25^, it will be interesting to determine if phospho-aspartate dependent EXO70D-mediated autophagic degradation also plays a role in the regulation of type-B ARRs as well.

## Methods

### Plant materials and growth conditions

All *Arabidopsis thaliana* lines used in this study are in the Col-0 ecotype. The GFP:ATG8f:HA autophagic marker lines were previously-described^79^. T-DNA insertion mutants of the *Exo70D* gene family (*exo70D1* (SALK_074650), *exo70D2* (WiscDsLox450H08), and the previously-described *exo70D3* (SAIL_175_D08)^48^ and *atg5-1* (CS39993)^61, 80^ were obtained from the *Arabidopsis* Biological Resources Centre, Ohio State University. Double (*exo70D1,2 exo70D1,3*, *exo70D1,2*) and triple (*exo70D1,2,3*) mutants were generated by crossing and confirmed by PCR-based genotyping using T-DNA and gene-specific primers (Table S1). TCSn::GFP;*exo70D1,2,3* was generated by crossing *exo70D1,2,3* to plants carrying a synthetic reporter for type-B ARR activity, *TCSn::*GFP^81^. To generate the ARR4:GFP, EXO70D3:GFP, mCherry:ATG8E and type-A ARR:CFP transgenic plants, each plasmid (see below) was introduced into *Agrobacterium* and transformed into Col-0 using floral dip^82^. The ARR4:GFP;*exo70D1,2,3,* ARR4:RFP;GFP:ATG8f, EXO70D3:GFP;mCherry:ATG8E, and ARR4:GFP;*atg5-1*lines were generated by crossing. All crosses were confirmed by antibiotic screening, PCR-based genotyping, and the presence of fluorescent tags by fluorescent microscopy.

For growth of *Arabidopsis* on plates, seeds were surface sterilized, plated on Murashige and Skoog (MS)^83^ media (MS basal salts, plus 1% sucrose and 0.8% Phytagel or phytoagar, pH. 5.8) and stratified at 4° C for 3 d. Unless otherwise stated, seedlings were grown for 10 days at 22° C under long day (LD) (16/8 h day/night). Adult plants were grown in soil at 22° C in either long-day (LD) (16 h light and 8 h dark) or short-day (SD) (8 h light and 16 h dark) photoperiod conditions as noted.

### Cloning and vector construction

CloneAmp HiFi PCR Premix (Takara) was used for all PCR amplifications, and the PCR products were gel-purified using GenCatch^TM^ Gel Extraction Kit (Epoch Life Sciences). Primers used for gene-cloning and genotyping are detailed in Table S1. Phospho-mimic (D-E) and phospho-dead (D-A) versions of the type-A ARRs were generated through site-directed mutagenesis of the conserved phosphorylation sites^37^. To create Entry Clones, blunt-end PCR products with directional overhangs were cloned into pENTR^TM^/D-TOPO^®^ vector using the pENTR^TM^ Directional TOPO^®^ Cloning Kit (Thermo Fisher Scientific). The resulting entry clones were inserted into respective Gateway-based destination vectors using Gateway^TM^ LR Clonase^TM^ II Enzyme Mix (Thermo Fisher Scientific). mCherry-ATG8E were generated by N terminal tagging of the full length ATG8E CDS sequence, using Greengate cloning^84^. The primers used for amplifying ATG8E CDS is given in Supplementary Table S1.

The CFP, GFP and RFP-tagged overexpression vectors were created by recombining the relevant entry clones into pUBC-CFP-Dest and pUBC-GFP-Dest and pUBC-RFP-Dest^85^, respectively. For mCherry-tagged vectors, the YFP cassette of pUBN-YFP-Dest was excised by *Spe*1 and *Mun1* restriction enzymes, and replaced with mCherry coding sequence^86^ flanked by *Spe*1 and *Mun1* restriction sites. mCherry_*Spe*1 and mCherry_Mun1 primers were used to introduce 5′ *Spe*1 and 3′ *Mun1* sites by PCR, and the fragments were purified after restriction digestion, mixed in an equimolar ratio and ligated using T4 DNA ligase (Fermentas).

Yeast two-hybrid vectors were generated by the Gateway^TM^-based recombination of respective Entry clones into the bait, pBMTN116c-D9, and prey, pACT2, destination vectors^87^. For bimolecular fluorescence complementation (BiFC), the type-A ARR and EXO70D3 Entry vectors were cloned into pUBN-cYFP-Dest and pUBN-nYFP-Dest^85^ destination vectors, respectively. For co-expression and co-immunoprecipitation assays in *N. benthamiana*, type-A ARRs, AHP2, full- and partial-length *EXO70D3* amplicons were cloned into myc-tagged pEarleyGate203 and HA-tagged pEarleyGate201 binary vectors^88^. Unless otherwise stated, destination clones were transformed into *Agrobacterium tumefaciens* strain, GV3101^89^, for *Arabidopsis* and *N. benthamiana* transformation. Whereas stable transgenic *Arabidopsis* plants were generated by *Agrobacterium*-based floral dip method^82^, genes were transiently expressed in *N. benthamiana* leaves by *Agrobacterium*-mediated leaf infiltration method^90^.

### Construction of phylogenetic tree and sequence alignment

The radial tree was created by the web-based Phylogeny^5, 6^ using Neighbor-joining algorithm. To generate the tree, the amino acid sequences were aligned using MUSCLE^TM7^, and curated using Gblocks^8, 9^ and the phylogeny created using PhyML^10^. Tree Rendering was performed using TreeDyn^11^. Bootstrap values represent 1000 iterations. Amino acid alignments were generated using T-Coffee^91^ and the web-based BoxShade software.

### Chemical treatment

For autophagic induction, seedlings were transferred in liquid MS (without sucrose) supplemented with 1 μM Concanamycin A (Conc A, Cayman Chemical, No. 11050)^92, 93^. Dimethyl sulfoxide (DMSO) served as vehicle control. For cytokinin treatment, seedlings were grown in MS media supplemented with sucrose and various concentrations of 6-benzylamino purine (BA, Sigma, No. B3408) or sodium hydroxide as control.

### Seedling cytokinin response assay

Responses to exogenous cytokinin was determined by a root elongation assay^37^. Briefly, surface-sterilized seeds were plated on vertical MS plates, stratified at 4° C for 3 days, and moved to growth chambers at 22° C, under 24 h light for 4 days. Seedlings were then transferred to fresh MS plates supplemented with NaOH as vehicle control or different concentrations of BA, and grown for 5 more days. Plates were scanned and root growth between days 4 and 9 measured using Fiji software^94^.

To determine the effect of cytokinin treatment on stability of ARR4:GFP, 10-day old seedlings were incubated in liquid MS medium supplemented with 5 μM BA for 0-12 h. Whole seedlings or roots were sampled and analyzed by immunoblot assay.

### Yeast two-hybrid analysis

Both bait and prey constructs were transformed into the L40ccαU strain^87^ using the previously-described lithium acetate transformation protocol^95^. Interactions were tested by the modified protocol of Dortey and co-workers^87^. Transformants were selected on -Leu-Trp Synthetic Complete (SC) double dropout media. Successful plasmid transformations were confirmed by colony PCR on the yeast colonies using gene-specific primers. To test interactions, cell suspensions of positive transformants were grown to an OD600 of 0.5, and 10 µL plated on -Leu-Trp-Ura-His media. Plates were incubated at 30° C for 4 days, and analyzed for interacting genes. Negative and Positive controls for each interaction included co-transforming with either the bait or prey vectors, and previously described interactors, respectively.

### Bimolecular Fluorescence Complementation Assay

*Agrobacterium* strain GV3101 carrying pUBN-cYFP-Dest-expressing EXO70D3 and phospho-mimic or phospho-dead ARR cloned into pUBN-nYFP-Dest were infiltrated into *N. benthamiana* epidermal leaf as previously described^90, 96^. Plasmids expressing the *tomato bushy stunt virus* suppressor p19^97^, was added to each infiltration mix to suppress host silencing of transgenes. The infiltrated *N. benthamiana* plants were grown for 3 days in constant light. Leaf discs from infiltrated plants were mounted in water and YFP and CFP signals were imaged in cells expressing both constructs using a Zeiss LSM710 confocal scanning microscope. To determine strength of interacting protein pairs, the fluorescent intensities of interacting proteins were quantified by Fiji software^94^.

### Co-immunoprecipitation assays

*Agrobacterium* strain GV3101 carrying plasmids expressing ARR5 and EXO70D3 was used to transiently transform leaf epidermal cells of *N. benthamiana* as described above. Infiltrated leaves were ground in liquid nitrogen, and total protein extracted in lysis buffer (1% glycerol; 25 mM Tris-HCl, pH 7.5; 1 mM EDTA; 150 mM NaCl; 10 mM DTT; 1 mM phenylmethylsulfonyl fluoride; 2% polyvinyl polyvinyl-polypyrolidone) supplemented with 1× protease inhibitor cocktail, cOmplete™ ULTRA Tablets, Mini, EDTA-free, EASYpack Protease Inhibitor Cocktail (Sigma). Co-immunoprecipitation was conducted using the μMACS™ Epitope Tag Protein Isolation Kits according to manufacturer’s instructions (Miltenyi Biotec), and purified on magnetic μ Column (Miltenyi Biotec). Columns were washed with wash buffer (1% glycerol; 25 mM Tris-HCl, pH 7.5; 1 mM EDTA; 250 mM NaCl; 10 mM DTT; cOmplete™ ULTRA Protease inhibitor Cocktail tablet), and proteins eluted with Elution Buffer (Miltenyi Biotec). Protein extracts were the subjected to immunoblot analyses using α-myc or α-HA antibodies.

### Immunoblot analyses

Proteins were resolved on SDS-PAGE followed by immunoblotting using α-HA high affinity antibody 3F10 (Sigma) and α-c-Myc Antibody (9E10): sc-40 (Santa Cruz Biotechnology) monoclonal primary antibodies, followed by goat anti-rat IgG–HRP:sc-2006 (Santa Cruz Biotechnology) or chicken anti-mouse IgG–HRP:sc-2954 (Santa Cruz Biotechnology) secondary antibody. For loading controls, membranes were probed with mouse α-tubulin sc-5286 (Santa Cruz Biotechnology) monoclonal primary antibody and anti-mouse secondary antibody. Signals were detected after incubating with SuperSignal™ West Femto Maximum Sensitivity Substrate (Thermo Fisher Scientific), and band intensities quantified by Fiji software^94^.

### Fluorescence microscopy analysis

Whole *Arabidopsis* roots and *N. benthamiana* leaf epidermal cells were imaged using a Zeiss LSM710 confocal scanning microscope equipped with 40 x 1.2 W C-Apochromat objective, an X-Cite 120 light-emitting diode fluorescent lamp and narrow-band fluorescent filter cubes. CFP, GFP, RFP and YFP were excited by 439-nm, 488-nm, 584-nm and 514-nm argon lasers, respectively, and collected by 485/20-nm, 516/20-nm, 610/10-nm and 540/25-nm nm emission filters. Fluorescent signals were detected using two HyD detectors in photon-counting mode (single sections) and normal mode (z-series). Bright-field images were taken using differential interference contrast optics and overlaid with fluorescence. To detect colocalization of autophagic vesicles, both fluorescence signals were sequentially line-captured using similar settings but a with narrower detection window. Images were collected and processed using ZEN 2009 software, and edited using Fiji software^94^. To calculate the colocalization of fluorescent signals, the images were thresholded using the Coste’s method^98^ and Manders’ Colocalization Coefficient (MCC)^47^ for overlays was calculated using Fiji software^94^.

### Quantitative Real Time PCR (qRT-PCR) analyses

Whole roots were isolated and total RNA extracted using the RNeasy Plus kit (QIAGEN). The RNA was DNase-treated using the RQ1 RNase-free DNase (Promega Corporation) and cDNA synthesized using Moloney Murine Leukemia Virus Reverse Transcriptase (M-MLV RT) and universal Random hexamers (Promega). qRT-PCR was performed with 2x PowerUP™ SYBR™Green Master Mix in the QuantStudio™ 6 Flex Real-Time PCR System (Applied Biosystems). *GAPDH* (AT1G13440) was used as housekeeping gene for normalization in all reactions. Primer sequences are provided in Table S1. Each sample was analyzed six times, including 3 biological replicates and 2 technical replicates each. Gene expression was determined using the ΔΔCt method of Pfaffl^99^ and presented as relative quantitation of target genes compared to the housekeeping genes.

## Acknowledgements

This work was supported by grant from the National Science Foundation (IOS-1856431) to JJK and (MCB 1856248) to JJK and GES.

## Author Contributions

AKA, CS, GES, and JJK conceptualized and designed research, with input from YD. AKA, CS and CYC conducted experiments. AKA and JJK wrote the manuscript with input from CS, GES and YD.

## FIGURE LEGENDS OF SUPPLEMENTARY MATERIALS

**Supplementary Figure 1: Effect of autophagy on protein and transcripts of ectopically expressed type-A ARR. (a)** Immunoblot assays of ectopically-expressed type-A ARRs in response to ConcA. Ten-day old *Arabidopsis* seedlings carrying *pUBQ10::ARR:CFP* constructs were treated for 8 h in sucrose-deficient liquid MS media supplemented with 1 µM ConcA (ConcA-8) or vehicle control (C-8). ARR protein quantities in treated roots, relative to untreated roots (Untrt), were analyzed by immunoblot assays using anti-GFP (α-GFP) antibodies. Anti-tubulin (α-tub) served as loading control. Rel. quantities represent ratio of intensity of α-GFP to α-tub band relative to ratio of Untrt bands for each blot. **(b)** qRT-PCR analyses of transcript levels of *pUBQ10::ARR:CFP* transgene in roots of seedlings of (a) above. **(c)** Immunoblot analyses of ectopically-expressed myc:ARR7 protein in roots of seedling in the presence of ConcA. myc:ARR7 levels were assayed using α-myc antibodies. **(d)** Representative confocal micrograph of the root elongation zones of *Arabidopsis* seedlings carrying CFP-tagged type-A ARR constructs. The light grown seedlings treated with ConcA (Upper panels) or DMSO as control (Lower panel) and exposed to carbon starvation for 18 h prior to imaging. Scale bar =10 µm. Quantification of the number of CFP-containing vacuolar vesicles. Data represents the average number of puncta per 100 µm^2^ of vacuole area, n=8. Statistical differences between treatments were analyzed using students’*t*-test; *p*<0.05. **** represent statistically different means. **(e)** qRT-PCR analyses of transcript of *pUBQ10::ARR4:GFP* transgene in roots of seedlings of wild-type and autophagy-deficient mutant, *atg5-1* seedlings treated with ConcA for 1 h. For (b) and (e), plots represent the mean normalized relative quantities (NRQ) of three biological replicates. Filled black circles and broken lines represent individual value and mean values, respectively. Statistical differences between genotypes was analyzed by Students’ *t*-test; *p*<0.05.

**Supplementary Figure 2: EXO70D3, a member of the EXO70 gene family, interacts with type-A ARRs in a phospho-Asp dependent manner. (a)** Phylogenetic tree showing the amino acid relatedness of 23 members of the EXO70 gene family. The genes cluster into three main clades, and nine subclades consisting of 3 EXO70As, 2 EXO70Bs, 2 EXO70Cs, 3 EXO70Ds, 2 EXO70Es and a single EXO70F1, 2 EXO70Gs, EXO700H1-EXO70H4, and EXO70H5-EXO70H8. The three isoforms of the EXO70D sub-clade are highlighted by grey halo. Bootstrap values represent 1000 iterations. Scale bar represents 0.2 amino acid substitutions per site. **(b)** Confocal micrograph of bimolecular fluorescence complementation (BiFC) assay depicting the interaction of EXO70D3 with the phospho-dead (D87A) and phospho-mimic (D87E) mutants of ARR5 in epidermal cells of *Nicotiana benthamiana* leaves. D87A and D87E mutations represent site-directed mutagenesis of the conserved Aspartate on the ARR5 to Alanine and Glutamate, respectively. EXO70D3 and ARR5 were tagged with N-terminus (nYFP) and C-terminus of split YFP (cYFP), respectively. NLS:CFP, a CFP-tagged nuclear localizing sequence was used as a infiltration control. Scale bar = 10µM. Quantification of the fluorescent intensities from images was performed with Fiji software. Data indicate the average of the maximum intensity values of the YFP signal relative to the corresponding CFP signals of 20 images for each infiltration. **** indicates statistically significant differences at *p*<0.001. **(c)** Confocal images showing EXO70D3:GFP protein predominantly localize in the cytoplasm of roots of four-day old *Arabidopsis* seedling. Seedlings carrying *pUBQ10::EXO70D3:GFP* construct were plasmolyzed by incubating for 90 min in MS media supplemented with 0.8 M mannitol and subsequently stained with propidium iodide for 1 min prior to imaging. Left panel indicates GFP channel, while middle and right panels represent propidium iodide (PI) and merged images, respectively. Plasmolyzed seedlings are shown in lower panel.

**Supplementary Figure 3: Full-length EXO70D3 destabilizes type-A ARRs *in planta*. (a)** A modified schematic representation of EXO70D3 as generated by the iLIR database^4^ and indicating ATG8-interacting motif (AIM) (232-LEWEVV-237) and an EXO70 domain (R^242^-D^609^). Three low complexity regions (S^59^-A^77^; A^148^-R^171^; S^518^-S^529^) were also identified (not indicated in Figure). In this study, interaction and co-expression assays were carried out using the AIM-containing EXO70D3^N-terminus^ fragment, and the rest of the protein, EXO70D3^C-terminus^. **(b)** Alignment of amino acid sequences of EXO70D isoforms. Yellow box and purple lines indicate AIMs and EXO70 domain, respectively. Black background indicates consensus residues. **(c)** Co-expressing phospho-mimic ARR5 (ARR5^D87E^) with full length EXO70D3 in leaves of *N. benthamiana* results in decrease in ARR5^D87E^ protein levels. HA:ARR5^D87E^ was co-infiltrated with equivalent amounts myc:EXO70D3 (left panel) or myc:AHP3 (right panel). **(d)** Full length EXO70D3 decreases ARR5 proteins in a concentration dependent manner. *N. benthamiana* leaves were transiently co-transformed with fixed amount of HA:ARR5 and increasing titer of myc:EXO70D3. **(e)** Co-expressing partial-length EXO70D3 does not result in the decrease in ARR5 proteins, regardless of the phosphorylation status of the conserved Aspartate. Phospho-mimic ARR5 (ARR5^D87E^) is co-expressed with N-terminus (EXO70D3^N-term^) (Left panel) and C-terminus (EXO70D3^C-term^) (Right panel) truncations of EXO70D3. **(f)** qRT-PCR analyses of transcript levels of *EXO70D1*, *EXO70D2* and *EXO70D3* in *Arabidopsis* roots and shoots. Plots represent the mean normalized relative quantities (NRQ) of three biological replicates with SEM. Statistical differences between values of root and shoot was analyzed by Student’ *t*-test, *P* < 0.05. Asterisk (*) indicate significant differences between root and shoot expression at *p=* 0.0139. **(g)-(h)** Effect of disrupting *EXO70Ds* on type-A ARR protein levels in *Arabidopsis*. Plants expressing *pUBQ10::ARR4:GFP* or *pUBQ10::ARR4:RFP* were crossed to *exo70D1,2,3* triple loss-of-function mutants to produce ARR4:GFP;*exo70D1,2,3* or ARR4:RFP;*exo70D1,2,3* plants. Confocal microscopy images indicating the expression of pUBQ10::ARR4:FP in whole root **(g)** and root tip **(h)** of wild-type (top panel) and *exo70D1,2,3* mutant (bottom panel) plants. Scale bars = 100 µm and 50 µm for (g) and (h), respectively. For **(h),** the signal intensity of individual nuclei were quantified. The graph represents the average signal measurement from 22 images of each genotype. Data was analyzed by unpaired Student’s t-test. **** indicate statistical difference at *p*=2.979×10^-030^

**Supplementary Figure 4: Gross morphology of mutants of *EXO70D* genes.** Comparative morphologies of 10-day and 6-week old wild type (Col-0) and *exo70D* mutants. Ten-day-old seedlings were grown under continuous light on vertical plates, while 6-week-old plants were grown under short day conditions in pots. Scale bar = 0.5 cm (for 10-day old seedlings); and 2 cm (for 6 weeks old

**Supplementary Figure 5: Interplay between cytokinin and EXO70Ds in *Arabidopsis* root. (a)** Effect of cytokinin on expression of *EXO70D* isoforms. qRT-PCR analyses of transcript levels of *EXO70D1*, *EXO70D2* and *EXO70D3* in *Arabidopsis* roots treated with BA or NaOH (Control). Plot represents NRQ values of expressions in BA-treated and Control samples from three biological replicates with two technical replicates each. Statistical differences between treatments was analyzed by unpaired Students’ *t*-test. **(b-d)** Effect of modulation of EXO70Ds on cytokinin responses in root. Effect of cytokinin on the length of primary roots of single (*exo70D1*, *exo70D2*, *exo70D3*) **(b)** and double (*exo70D1,2*, *exo70D1,3*, *exo70D2,3*) **(c)** loss-of-function mutants of *EXO70D* genes. Seedlings were grown on vertical MS media and after 4 days transferred to plates supplemented with 6-benzyl-adenine (BA) or NaOH plates as a vehicle control for 6 days. Graphical representation of quantitation of the primary root length in response to different BA concentrations. Values represent the average of more than 11 determinations, analyzed with one-way ANOVA followed by Tukey-Kramer multiple mean comparisons at *p<*0.05. **(d)** Disrupting all three *EXO70D* genes reduces the type-B ARR activity in *Arabidopsis* roots. Confocal microscopy images showing expression of the type-B RR reporter TCSn::GFP in wild type (left panels) and *exo70D1,2,3* triple mutant (right panels) plants incubated in 5µM BA (lower panels) or NaOH control media (upper panels) for 2h. Scale bar = 50 µm. Intensity of the GFP signal from 20 determinations (n=20) was analyzed with one-way ANOVA and Tukey-Kramer comparisons; *P* < 0.05. Asterisk (*) indicate significant differences at indicated *P*-values.

**Supplementary Figure 6: Responses of lower order mutants of *EXO70D*s to fixed-carbon starvation.** Seedlings were grown for six weeks on potted soil under short-day conditions, and transferred to dark for 7, 9, 11, 13 days. Plants were recovered in the light for 7 days. **(a)** Representative images of single (*exo70D1, exo70D2, exo70D3*) and double (*exo70D1,2*; *exo70D1,3*; *exo70D2,3*) loss-of-function mutants of *EXO70D*s following 13-day-dark treatment. Col-0 and *atg5-1* served as wild-type and autophagy deficient controls, respectively. Graphical representation of quantification of survival of single **(b)** and double **(c)** mutants in response to carbon starvation. Survival was estimated as the percentage of plants with new leaves after the dark treatment. Values represent mean ± SEM percentage survival of 3 biological replicates. Each biological replicate consisted of 8 plants per genotype per treatment.

