## Supplementary Fig. 1 for "EXO70D isoforms mediate selective autophagic degradation of Type-A ARR proteins to regulate cytokinin sensitivity"

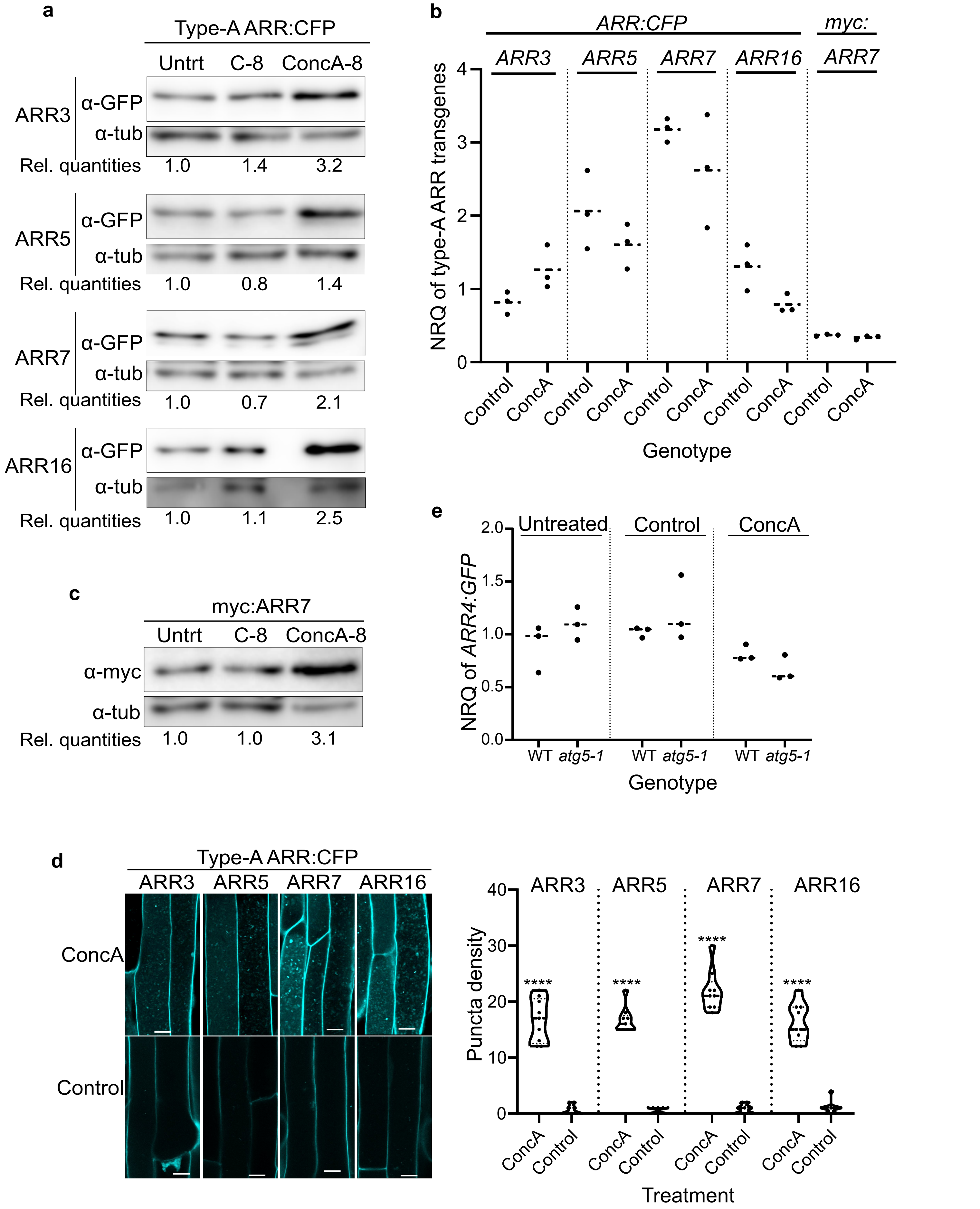


**Supplementary Fig. 1:** Effect of autophagy on protein and transcripts of ectopically-expressed type-A ARRs
