## Supplementary Fig. 2 for "EXO70D isoforms mediate selective autophagic degradation of Type-A ARR proteins to regulate cytokinin sensitivity"

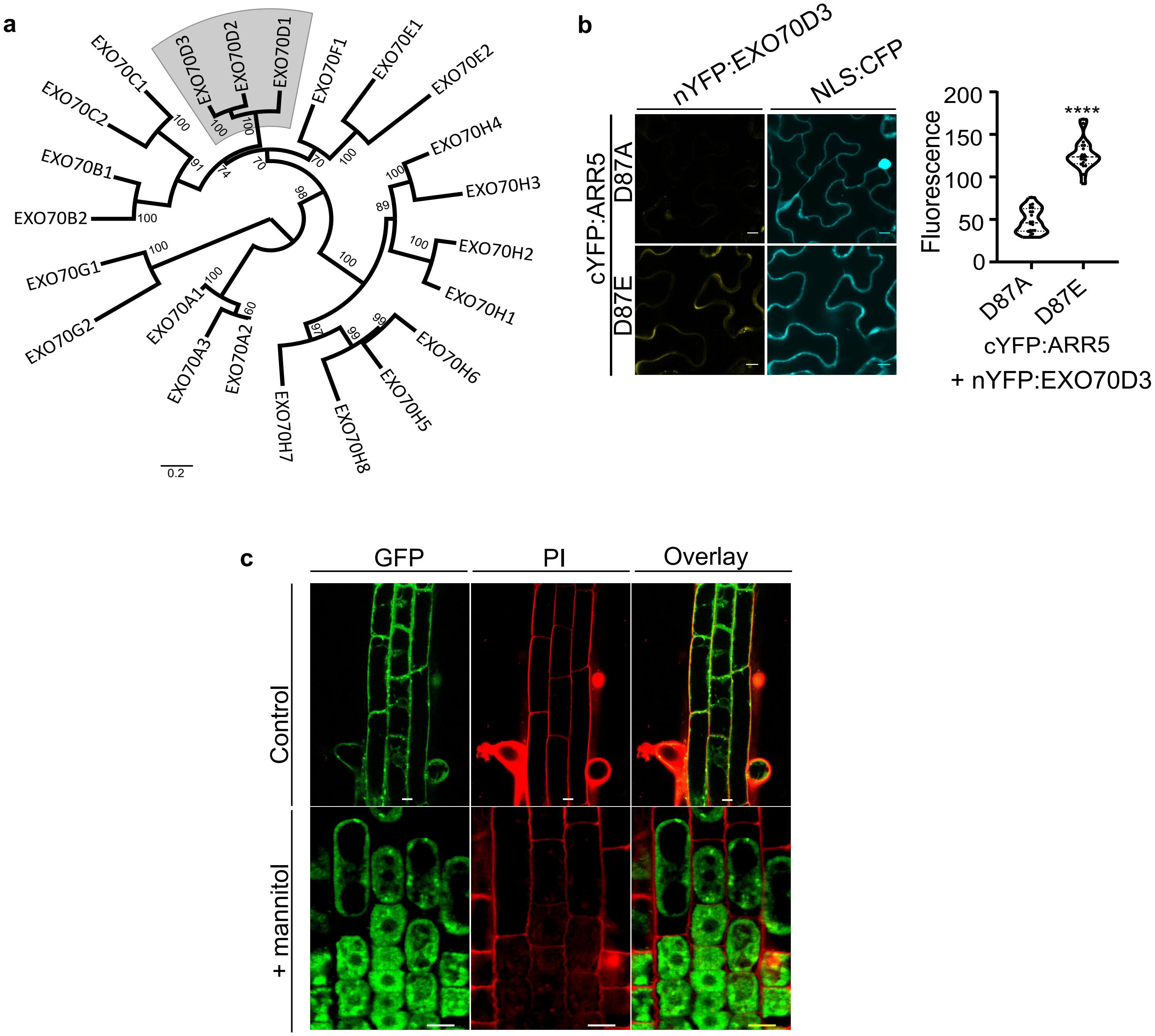


**Supplementary Fig. 2:** EXO70D3, a member of the EXO70 gene family, interacts with type-A ARRs in a phospho-Asp dependent manner
