## Supplementary figures and images for "EXO70D isoforms mediate selective autophagic degradation of Type-A ARR proteins to regulate cytokinin sensitivity"

### Supplementary Fig. 3

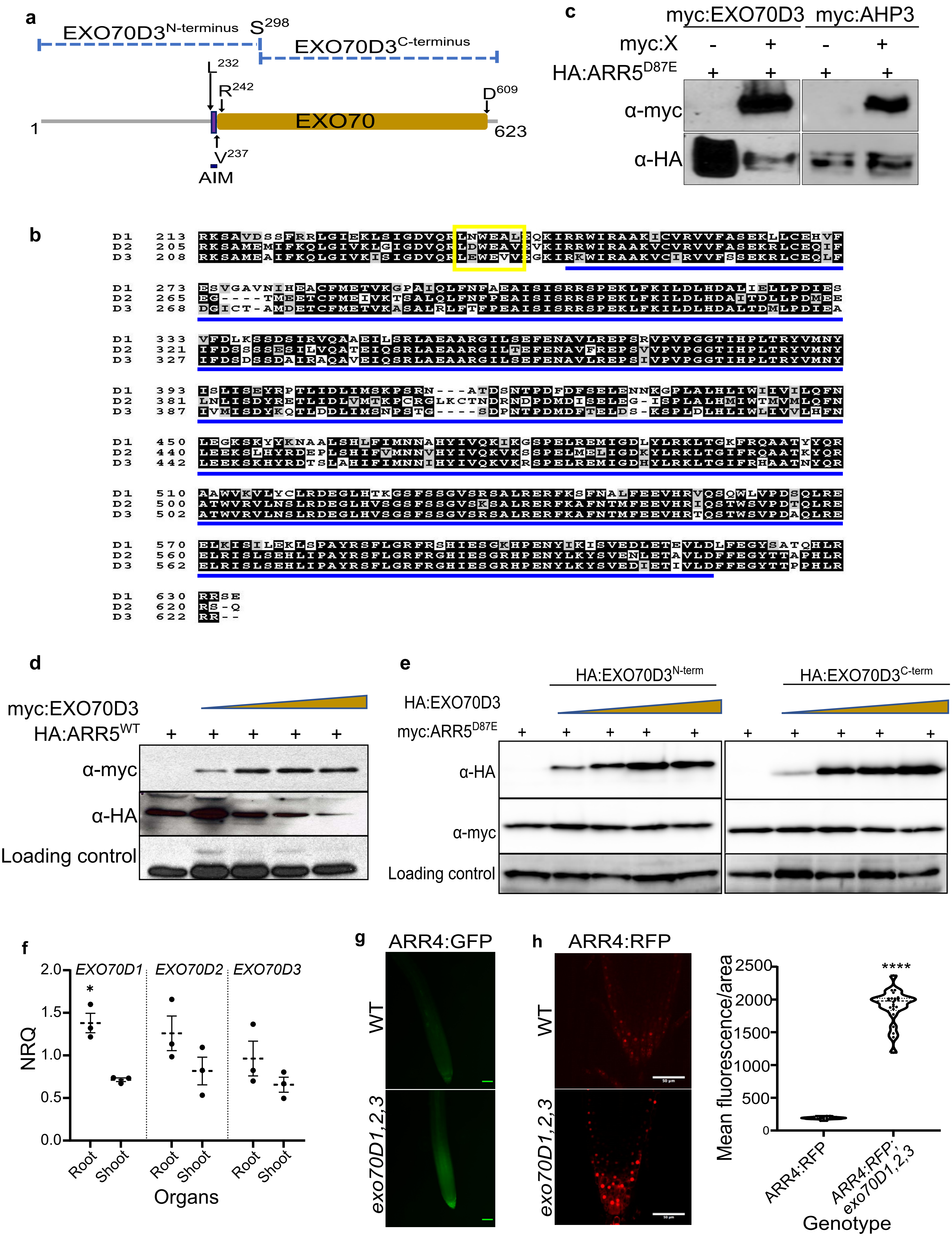


**Supplementary Fig. 3:** Full-length EXO70D3 destabilizes type-A ARR proteins *in planta*

### Supplementary Fig. 4

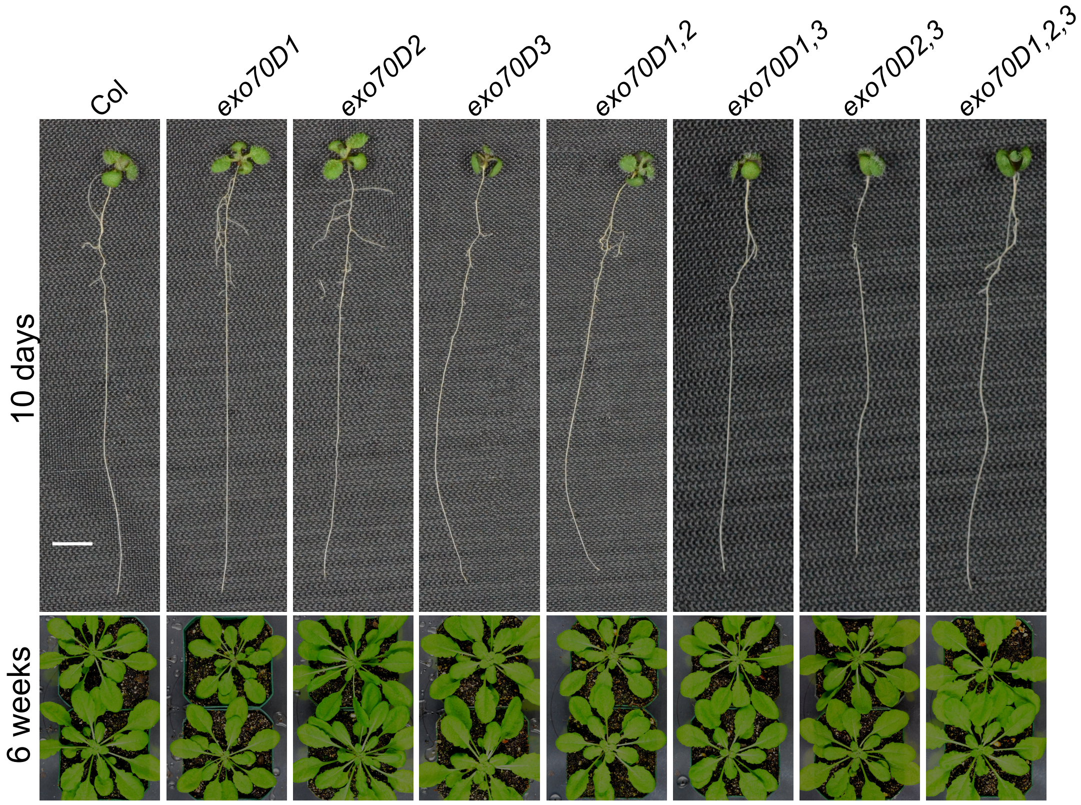


**Supplementary Fig. 4**: Morphology of mutants of *EXO70D* genes

### Supplementary Fig. 5

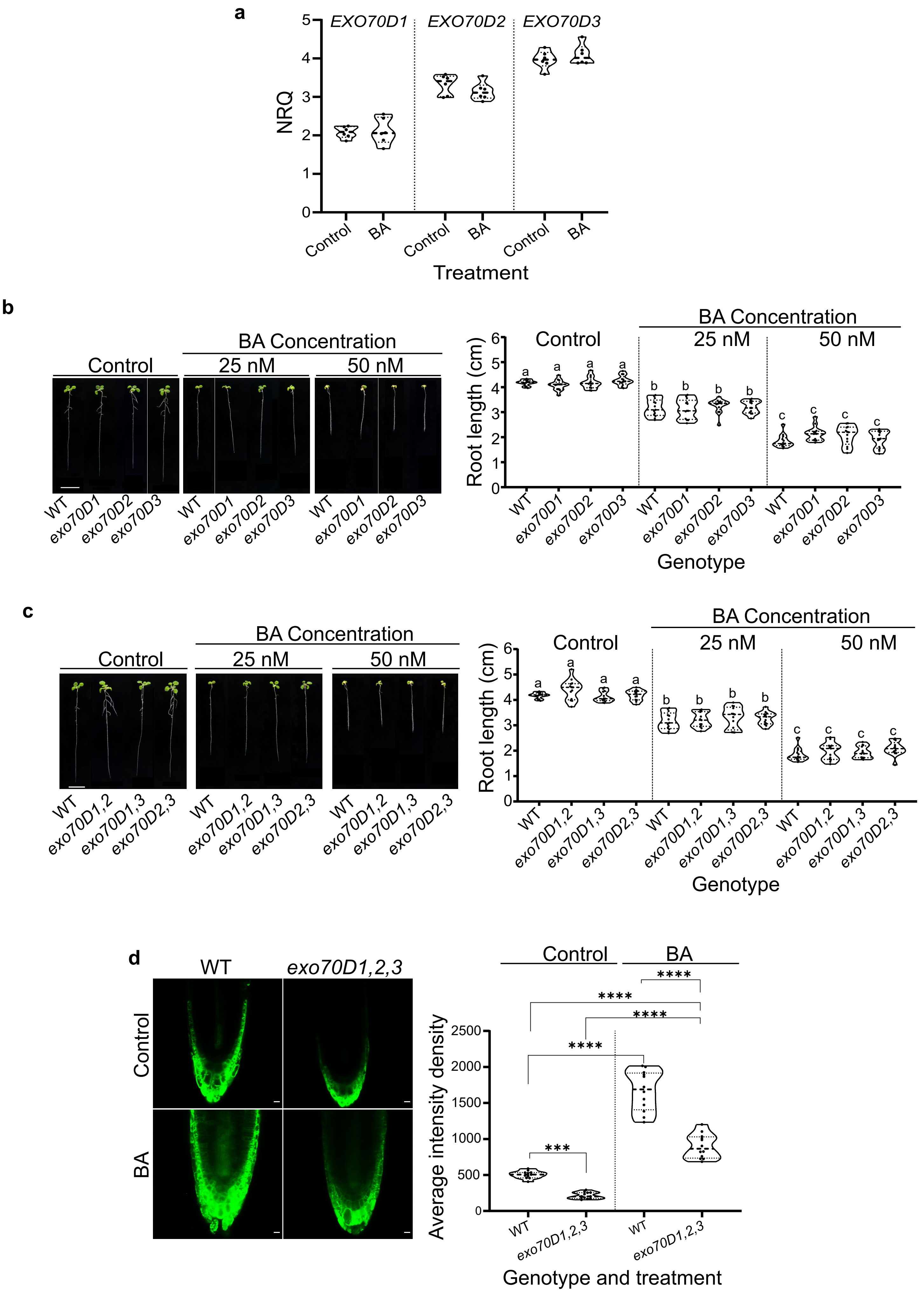


**Supplementary Fig. 5**: Interplay between cytokinin and EXO70Ds in Arabidopsis root
