## Supplementary Fig. 6 for "EXO70D isoforms mediate selective autophagic degradation of Type-A ARR proteins to regulate cytokinin sensitivity"

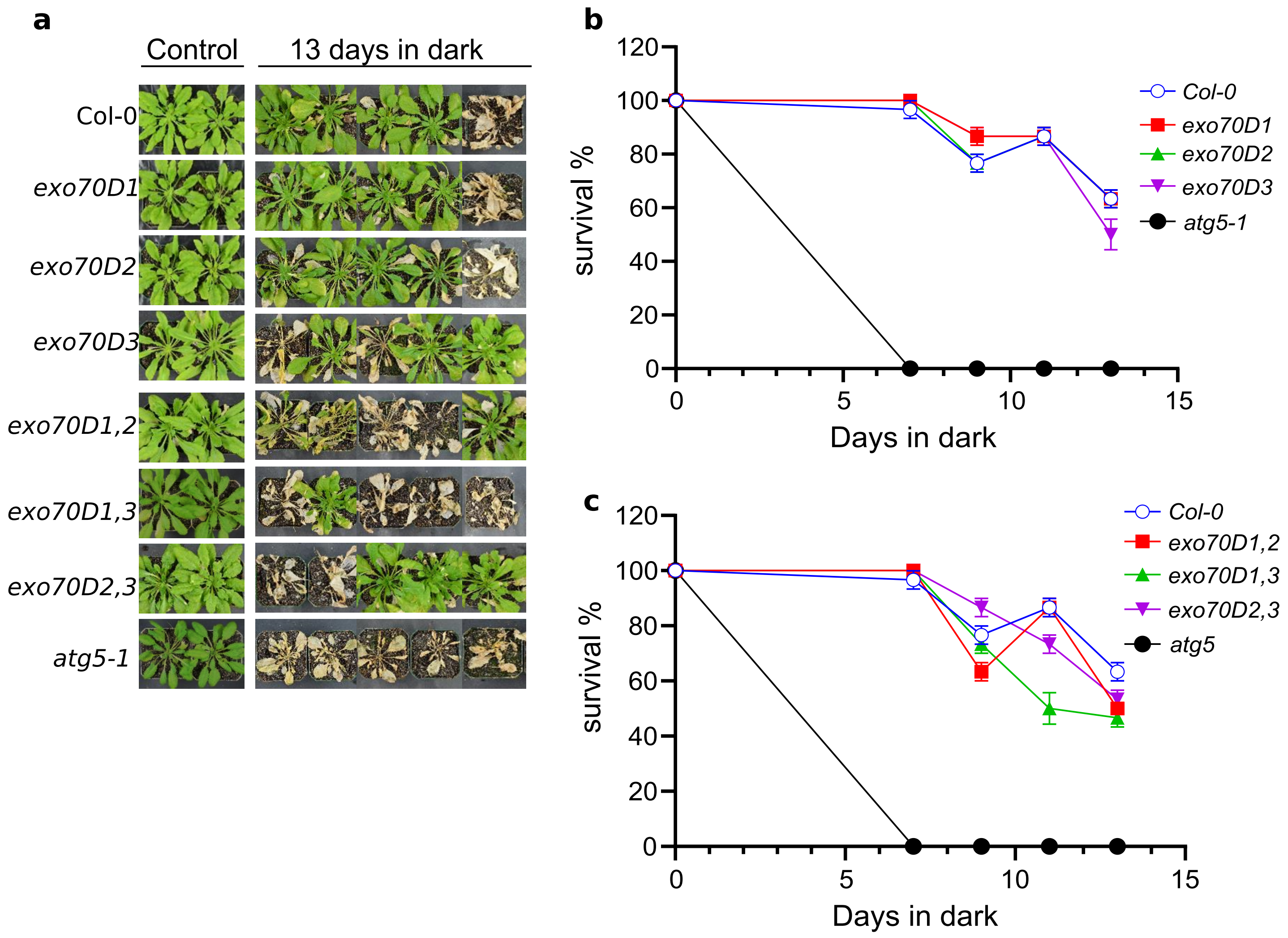


**Supplementary Figure 6:** EXO70Ds and type-A ARRs modulate plant response to fixed carbon starvation
