## Supplementary material for "EXO70D isoforms mediate selective autophagic degradation of Type-A ARR proteins to regulate cytokinin sensitivity": Table S1

**Table S1: Primers used in this study**

| **Gene** | **Forward primer (5’→3’)** | **Reverse primer (5’→3’)** | **Purpose** |
| --- | --- | --- | --- |
| *exo70D1* | ggatcgtcttctcagatgctg | tgctctgtgagcatgtttttg | T-DNA genotyping |
| *exo70D2* | cgcattctcaaactcggtaag | attctgattatggcaacaccg | T-DNA genotyping |
| *exo70D3* | cgtctttaaccgtcgaagttg | ctgacccatgtagctctctgg | T-DNA genotyping |
| *atg5-1* | atttgctatttgtttggcacg | taccgttcatgacagaggtcc | T-DNA genotyping |
| *arr3* | ggaactagtagcaatatctctcttctatcttttc | cacagaggtaaactgtcacacattatttg | T-DNA genotyping |
| *arr4* | tttatgtgcgacacgttgatgactacttt | ggaggcgcgagagattaaagggacatctat | T-DNA genotyping |
| *arr5* | tctctctgtggtacatttcttgaaaaatggg | cttggggaaatttctaagaaaagccatgta | T-DNA genotyping |
| *arr6* | tgtagaagttaaatgcgtgaacttccaca | gctatggtgaatcctcttgacaagttactc | T-DNA genotyping |
| *arr7* | ggcggtttgcagactcacttacctga | gactctctcaaacattgtcttt | T-DNA genotyping |
| *arr8* | caaatggctgttaaaacccaccaata | ccattgttagtgtgctatcacctgagtg | T-DNA genotyping |
| *arr9* | ggatcccagactctttatttctcttcctc | cccacatacaacatcatcatcatattcc | T-DNA genotyping |
| *arr15* | ccatttattctcctctcatctc | atctaatcatccccatctcc | T-DNA genotyping |
| *EXO70D1* | caccatggaaccacatgaccaaactcacg | tcactcggatcgtcttctcagatgc | Cloning full-length CDS |
| *EXO70D2* | caccatggcaacaccggagact | tcactgagaccgtctcaaa | Cloning full-length CDS |
| *EXO70D3* | caccatggaaccgccggagaat | ttatcgcctcctcaagtgtggt | Cloning full-length CDS |
| *ATG8a* | caccatgatctttgcttgcttga | tcaagcaacggtaagagatc | Cloning full-length CDS |
| *ATG8b* | caccatggagaagaactccttca | ttagcagtagaaagatccac | Cloning full-length CDS |
| *ATG8c* | caccatggctaatagctctttca | ctaagaagttgtgtgtttac | Cloning full-length CDS |
| *ATG8d* | caccatggcgattagctccttca | ttagaagaagatcccgaacg | Cloning full-length CDS |
| *ATG8e* | caccatgaataaaggaagcatct | ttagattgaagaagcaccga | Cloning full-length CDS |
| *ATG8f* | caccatggcaaaaagctcgttcaa | ttatggagatccaaatccaa | Cloning full-length CDS |
| *ATG8g* | caccatgagtaacgtcagcttcag | ttaagtcattgacgatccaa | Cloning full-length CDS |
| *ATG8h* | caccatggggattgttgtcaagt | ttagccgaaagttttctcgg | Cloning full-length CDS |
| *ATG8i* | caccatgaaatcgttcaaggaac | tcaaccaaaggttttctcac | Cloning full-length CDS |
| *mCherry:ATG8e* | aacaggtctcaggctctatgaata  aaggaagcatctttaagatg | aacaggtctcactgagattgaa  gaagcaccgaatgt | Cloning full-length CDS for Greengate cloning |
| *EXO70D3^N-term^* | caccatggaaccgccggagaat | ttaacttatagcttcaggg | Cloning CDS |
| *EXO70D3^C-term^* | caccatgatcagtagaagatcacctg | ttatcgcctcctcaagtgtggt | Cloning CDS |
| *AHP2* | caccatggacgctctcattgctca | ttagttaatatccacttgagg | Cloning full-length CDS |
| *AHP3* | caccatggacacactcattgctca | ttatatatccacttgagg | Cloning full-length CDS |
| *ARR4* | caccatggccagagacggtggt | ctaatctaatccgggactcc | Cloning full-length CDS |
| *ARR5* | caccatggctgaggttttgcgtc | ttagatctttgcgcgttttag | Cloning full-length CDS |
| *ARR7* | caccatggcggttggtgaggt | tcaaagtagagaaaaaaggtt | Cloning full-length CDS |
| *ARR16* | caccatgaacagttcaggaggttc | ttagcttctgcagttcatga | Cloning full-length CDS |
| mCherry_Spe1_F | tccaactagtatggtgagcaagggcgagg |  | Cloning full-length CDS with *Spe1* and *Mun1* sites |
| mCherry_Mun1_R |  | gattaacaattgcttgtacagctcgtcca | Cloning full-length CDS with *Spe1* and *Mun1* sites |
| *EXO70D1* | aatggtctccgccggttatc | agcgcttcccaatttagcct | qRT-PCR |
| *EXO70D2* | agactcgtggcgttgattcc | cttctcgatcaccaccgctt | qRT-PCR |
| *EXO70D3* | ggaaccgccggagaatagtt | atccacctcatcacgatcgc | qRT-PCR |
| *ARR3:FP* | tggttgacgatgaagattcg | accactttgtacaagaaagctgggt | qRT-PCR of transgene |
| *ARR4:FP* | gaagatgatgacgtgttgacg | accactttgtacaagaaagctgggt | qRT-PCR of transgene |
| *ARR5:FP* | ttaatgaaagctgaggaaagagc | accactttgtacaagaaagctgggt | qRT-PCR of transgene |
| *ARR6:FP* | tcacaaagagtattcagaagagagagc | accactttgtacaagaaagctgggt | qRT-PCR of transgene |
| *ARR7:FP* | gagaaagcttcaagaagacagtga | accactttgtacaagaaagctgggt | qRT-PCR of transgene |
| *ARR9:FP* | aacgttttcattgtgtcttgattc | accactttgtacaagaaagctgggt | qRT-PCR of transgene |
| *ARR16:FP* | agcgagtggagctcaaatgt | accactttgtacaagaaagctgggt | qRT-PCR of transgene |
| *GAPDH* | aggccatcaaggaggaatct | gaaaatgcttgacctgttgtcac | qRT-PCR |
